# Peptidome Surveillance Across Evolving SARS-CoV-2 Lineages Reveals HLA Binding Conservation in Nucleocapsid Among Variants With Most Potential for T-Cell Epitope Loss In Spike

**DOI:** 10.1101/2022.03.18.484954

**Authors:** Kamil Wnuk, Jeremi Sudol, Patricia Spilman, Patrick Soon-Shiong

**Author notes:** Corresponding author: Kamil Wnuk.

## Abstract

To provide a unique global view of the relative potential for evasion of CD8+ and CD4+ T cells by SARS-CoV-2 lineages as they evolve over time, we performed a comprehensive analysis of predicted HLA-I and HLA-II binding peptides in spike (S) and nucleocapsid (N) protein sequences of all available SARS-CoV-2 genomes as provided by NIH NCBI at a bi-monthly interval between March and December of 2021. A data supplement of all B.1.1.529 (Omicron) genomes from GISAID in early December was also used to capture the rapidly spreading variant. A key finding is that throughout continued viral evolution and increasing rates of mutations occurring at T-cell epitope hotspots, protein instances with worst case binding loss did not become the most frequent for any Variant of Concern (VOC) or Variant of Interest (VOI) lineage; suggesting T-cell evasion is not likely to be a dominant evolutionary pressure on SARS-CoV-2. We also determined that throughout the course of the pandemic in 2021, there remained a relatively steady ratio of viral variants that exhibit conservation of epitopes in the N protein, despite significant potential for epitope loss in S relative to other lineages. We further localized conserved regions in N with high epitope yield potential, and illustrated HLA-I binding heterogeneity across the S protein consistent with empirical observations. Although Omicron’s high volume of mutations caused it to exhibit more epitope loss potential than most frequently observed versions of proteins in almost all other VOCs, epitope candidates across its most frequent N proteins were still largely conserved. This analysis adds to the body of evidence suggesting that N may have merit as an additional antigen to elicit immune responses to vaccination with increased potential to provide sustained protection against COVID-19 disease in the face of emerging variants.

## Introduction

The ability to understand and predict the potential of new SARS-CoV-2 variants to escape immune responses elicited by previous lineages is vital to meeting and overcoming the challenge of the COVID-19 pandemic. Amidst the multitude of parallel efforts to characterize the adaptive immune response to the virus (1–3) and the potential for it to be sustained against variants, many have highlighted T-cell responses as playing an important role in disease control, resolution, and immunity (4–10). Underscoring the importance of T cells, earlier reports revealed that CD4+ and CD8+ T-cell responses against the spike (S), membrane (M), and nucleocapsid (N) proteins of original SARS-CoV persisted for up to 11 years post infection (11) and N-specific T-cell reactivity to 17 years post infection (12).

Persistence of T-cell reactivity against viral proteins does not, however, indicate that such pre-existing cellular immunity will be sufficiently protective against highly mutated viral variants. This risk has come to the forefront in the present COVID-19 pandemic because SARS-CoV-2 variants more transmissible than the original strain have emerged at a rapid rate and spread throughout the globe.

There is some encouraging evidence that, due to the broad CD4+ and CD8+ T-cell responses against multiple SARS-CoV-2 structural proteins (7, 10, 13–15), it is unlikely that variants will escape the majority of T-cell immunity conferred by prior infection or vaccination. In a study of peripheral blood mononuclear cells (PBMCs) from 57 recovered or vaccinated subjects, Tarke *et al*. (16) reported that on average 93% of CD4+ and 97% of CD8+ T-cell epitopes verified from the reference SARS-CoV-2 genome (14) were conserved across genomes selected to represent 4 Variants of Concern (VOC). Across the 4 variants, these conserved epitopes were found to account for an average of 91.5% and 98.1% of CD4+ and CD8+ T-cell recognition, respectively. The authors went on to demonstrate that COVID-19 patients showed no significant difference in T-cell reactivity to peptide pools spanning 4 VOC lineages compared to peptides sourced from the SARS-CoV-2 reference/ancestral genome (NCBI Reference Sequence: NC_045512). This was true not only for peptide pools specific to S, but also those spanning proteins across the whole viral genome. Individuals who had received mRNA vaccines showed some diminished T-cell response to S B.1.351 and B.1.427/B.1.429 variant peptide pools, but overall, responses were considered robust. Conversely, Lucas *et al*. (17) found decreased CD8+ (but not CD4+) T-cell responses against S peptides from the P.1 lineage compared to peptides from the reference/ancestral lineage in individuals who had received mRNA vaccines. Further, in recovered COVID-19 patients, Agerer *et al*. (18) confirmed that single mutations in CD8+ T-cell epitopes led to diminished HLA-I binding, a weaker T-cell response, and ineffective cytotoxicity in HLA-matched COVID-19 patients. Thus, mutations in SARS-CoV-2 may evade aspects of the adaptive immune response at the narrow scope of specific epitopes and HLAs.

Additionally, individuals or populations with significantly different HLA phenotypes than those present in the referenced study cohorts may exhibit different levels of evasion risk. This may be one possible explanation for conflicting findings across studies (16, 17). As another example, even within a geographically limited cohort, CD8+ T-cell epitopes were found to vary significantly based on patient HLA-I repertoires (14).

Even if the risk for T-cell evasion by variants has not, thus far, appeared to be great, given the emergence of highly-mutated new variants such as B.1.1.529 (Omicron) (19), continued monitoring of the potential impact on T-cell reactivity is warranted. Our approach to understanding the implications of viral evolution and provide a comprehensive view of CD4+ and CD8+ evasion potential is the assessment of the dynamics of epitope – and specifically predicted HLA binding - changes with time. HLA presentation is an essential precursor to T-cell recognition, and recent results have demonstrated that immunodominant CD4+ T-cell epitopes specific to SARS-CoV-2 correlate with HLA binding promiscuity (14). We analyzed both the S and N peptidomes, motivated by next generation vaccines currently in development that include both S and N antigens (20–22) and findings discussed above indicating broad T-cell response across structural proteins.

We identified viral protein regions predicted to be the highest frequency sources of HLA-I and HLA-II binding peptides, and tracked the evolution of binding loss (and thus latent potential for T-cell evasion) across all available viral genomes at bi-monthly time points throughout 2021. Narrowing our focus to specific protein locations with peak frequencies of potential epitopes, that is, potential epitope hotspots, improved robustness to noise when ranking viral variants according to risk of epitope loss, and simplified identification of critical regions or HLAs most impacted by epitope loss.

Our predictive analysis differs from that of others by focusing on the evolution of SARS-CoV-2 viral lineages, integrating results across a broad range of peptide lengths, considering impacts of observed mutation co-occurrence by analyzing full protein sequences versus independently considering key mutations, demonstrating efficacy of CD4+ T-cell epitope predictions, and by offering a distinct approach to capturing a comprehensive set of HLA-I and HLA-II alleles. In an effort to address fairness in studying the potential impact of SARS-CoV-2 variants on world populations, we sampled a representative set of HLA-I and HLA-II alleles based not only on documented frequency of appearance, but also leveraged learned embeddings to capture functionally similar HLA clusters. These functional groupings were then used in the hope of ensuring that a uniquely behaving set of HLAs was not excluded due to absence in majority populations.

Access to functional embeddings was enabled by in-house neural networks trained to predict peptide binding to HLA-I and HLA-II molecules. In our tests, the models compared favorably to the state-of-the-art (NetMHCpan-4.1), and opened the door to additional capabilities including abstaining from classification in cases of ambiguity, and the ability to visualize and leverage intermediate representations of HLAs or peptides.

Over the time period analyzed, there remained a relatively steady ratio of viral variants that exhibited conservation of N epitopes among lineages exhibiting most potential for epitope loss in S. VOC lineages generally had greater overall loss in binding for both HLA-I and HLA-II than VOI lineages across both S and N proteins. This correlates with the expectation of increased potential for immune evasive viral variants to emerge over time as the pandemic progresses, highlights that lineages exhibiting more rapid spread also have more opportunity to develop evasion, and underscores the critical role of genomic surveillance to uncovering how the immune landscape of viral variants may be shifting. Interestingly, we also found that within evolutionary trajectories of VOC and VOI lineages, protein instances exhibiting worst case binding loss have not become the most frequent versions of S or N for any lineage; which suggests T-cell evasion may not be a dominant evolutionary pressure on SARS-CoV-2. This observation held true throughout the initial surge of the Omicron variant, which despite its high rate of mutations in S compared to other VOCs, also demonstrated epitope conservation in the N protein. Finally, we used our analyses to identify conserved regions in N most likely to yield immunodominant CD4+ and CD8+ T-cell epitopes, and illustrate heterogeneity of CD8+ T-cell response in the S protein across our comprehensive set of HLAs.

We provide our snapshots of the evolution of the SARS-CoV-2 peptidome as interactive visualizations at: https://research.immunitybio.com/scov2_epitopes/. The interactive visualization link also includes in-depth examples demonstrating localization of potential epitope loss for most frequent versions of S and N proteins corresponding to VOC lineages.

## Results

### Localization of potential epitope hotspots on viral proteins

We observe that when HLA binding predictions are evaluated at all possible peptides in a sliding window along a source protein (see *Methods*), certain regions exhibit an increased frequency of high confidence predictions relative to elsewhere on the same protein. This observation was leveraged to narrow the search space of our analysis only to protein regions most likely to impact CD8+ or CD4+ T-cell responses. Potential epitope hotspot selection serves to improve robustness to noise (from protein locations less likely to be impactful to immune response) when globally comparing viral variants, and enables quick localization of where potential epitope loss is most significant when more resolution is desired.

For each protein considered, hotspot regions were identified for a representative set of HLA-I only, HLA-II only, and an aggregate that integrated all HLA-I and HLA-II binding prediction information to create a map of pan-HLA potential epitope hotspots. Representative sets of HLA-I and HLA-II alleles used to inform hotspot selection were chosen by combining both global allele frequency information as well as functional clustering based on embeddings from our trained HLA binding prediction models, to ensure rare but functionally unique HLAs were represented. Potential epitope hotspot regions are shown in **Figure 1** for six critical SARS-CoV-2 proteins in the reference genome: S, N, membrane (M), envelope (E), open reading frame 1ab (ORF1ab) and ORF3a of NCBI Reference Sequence NC_045512 (23).

**Figure 1.**
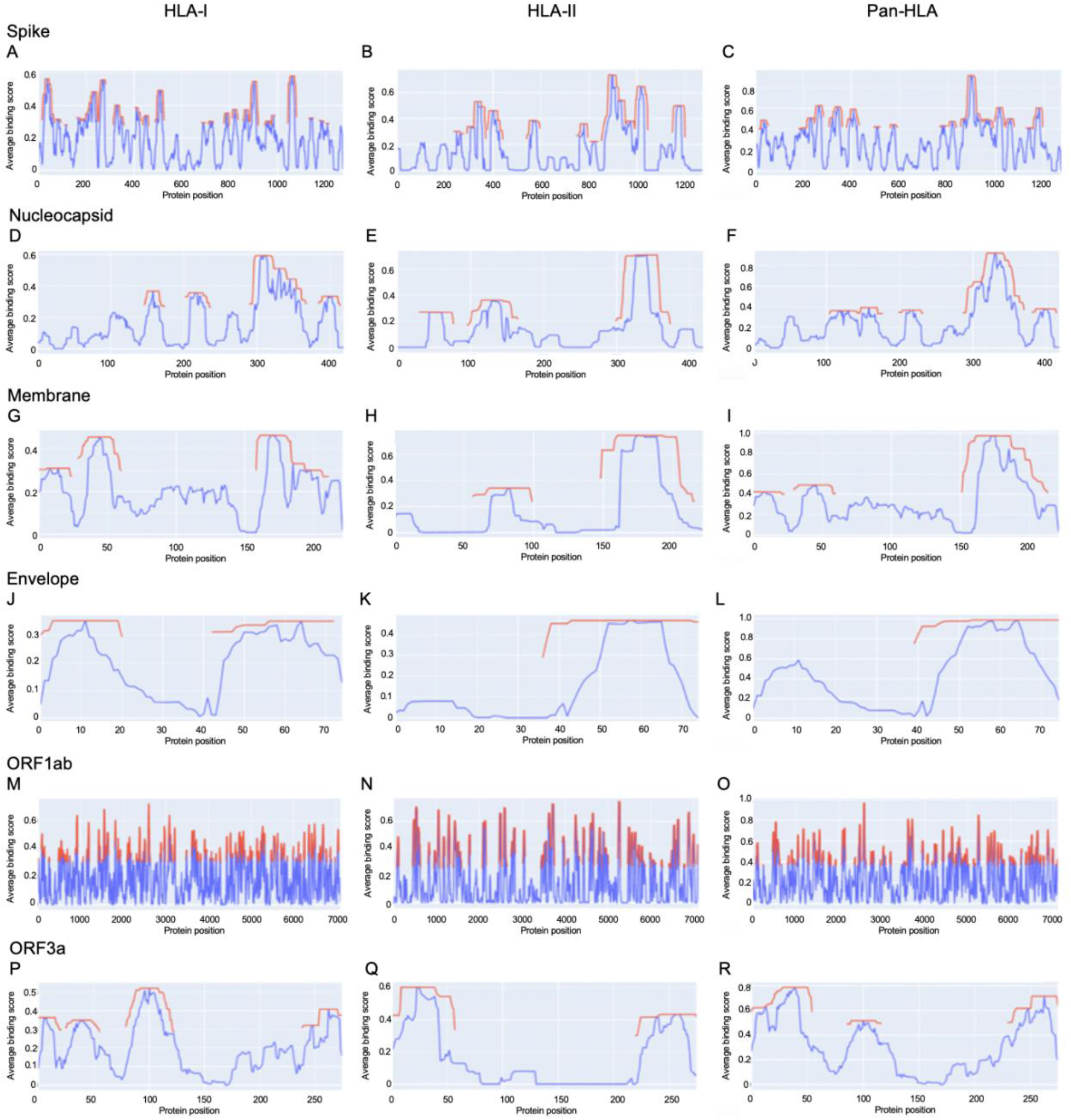
Epitope hotspots in six SARS-CoV-2 proteins. Protein regions with peak frequency of predicted binding peptides (potential epitope hotspots) across HLAs are indicated in red for 6 key proteins from the SARS-CoV-2 reference genome (NCBI Reference Sequence: NC_045512): spike (A-C), nucleocapsid (D-F), membrane (G-I), envelope (J-L), open reading frame 1ab (ORF1ab) (M-O), and ORF3a (P-R). Red lines show the value of the nearest maxima of the aggregate signal (blue) within a set sliding window size (9 amino acids for HLA-I, 15 for HLA-II, 12 for pan-HLA). For each protein we show hotspots on aggregate signals across all HLA-I molecules only (left column), HLA-II only (middle), as well as the combined pan-HLA signal (right).

S was found to have 18 HLA-I binding hotspots, 8 HLA-II binding hotspots, and 12 pan-HLA binding hotspots (**Fig. 1A-C**); whereas the much shorter N protein featured 4 HLA-I hotspots, 3 HLA-II hotspots, and 4 pan-HLA hotspots (**Fig. 1D-F**).

Several notable locations on the S protein fall within our hotspot locations. For example, the receptor binding domain (RBD) of S that interfaces with host cell angiotensin-converting enzyme 2 (ACE2) (24) overlaps pan-HLA hotspots 4, 5; HLA-I hotspots 6, 7, 8; and HLA-II hotspot 3. From among all the Centers for Disease Control (CDC) listed substitutions of therapeutic concern (implicated in contributing to increased transmissibility, severity, or reduced susceptibility to therapies) based on data up to July 31, 2021 (25); only E484K does not fall within any hotspots. For all others: L452R falls within HLA-I hotspot 7; K417N/K417T is in pan-HLA hotspot 8, HLA-I hotspot 6, HLA-II hotspot 3; and N501Y falls in pan-HLA hotspot 5 and HLA-I hotspot 8.

In alignment with our analysis, response frequencies from empirically confirmed epitopes aggregated across 25 studies spanning 1197 human subjects (9) aligned well with our predictions (**Fig. 2; Table 1**). Further, epitopes verified at the scope of individual studies such as by Saini et al. (15) were shown to consistently be in zones that align with our hotspot predictions, and were consistently assigned high HLA binding confidence values (**Tables 2 and 3**).

**Figure 2.**
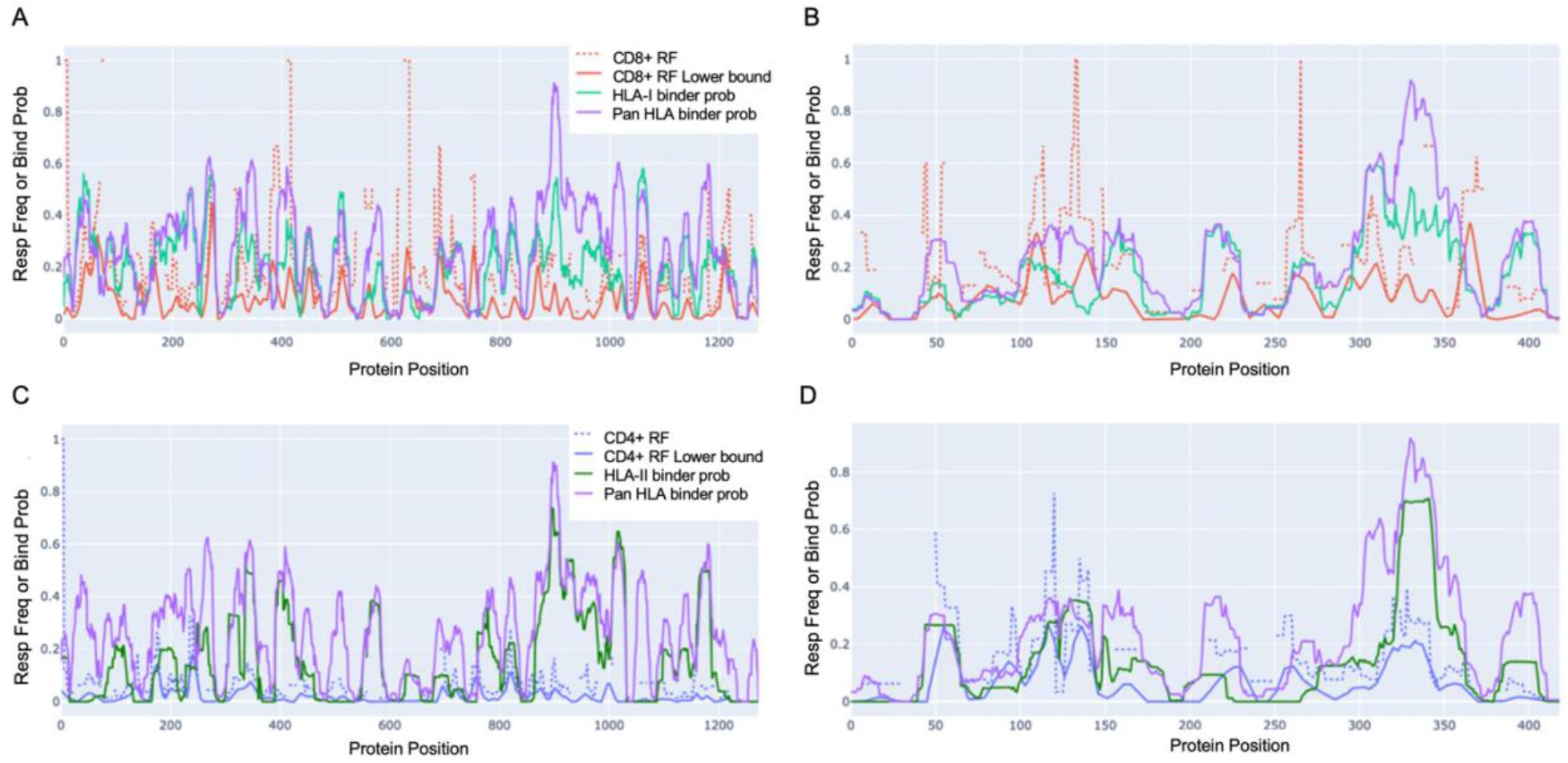
Epitope response frequency correlates with aggregated binding predictions in SARS-CoV-2 proteins. For CD8+ T-cell epitopes collected from across 25 studies by Grifoni et al. (9) we found position-specific response frequency (RF) (dash red) and RF lower bound (95% confidence interval) averaged with a 10 amino acid sliding window (solid red) correlated with our aggregated HLA-I (turquoise) and pan-HLA (purple) binding prediction scores in the S (A) and N (B) proteins. CD4+ T-cell epitope RF (dash blue) and RF lower bound (solid blue) also correlated with HLA-II (green) and pan-HLA aggregated predictions for S (C) and N (D). Legend in A applies to B; legend in C applies to D.

**Table 1.** Correlation and intersection at peaks of epitope response frequency with aggregated HLA binding predictions.

| Protein | T-cell epitope RF restriction | Aggregate HLA prediction class | Spearman $r_s$ | Fraction of RF peaks that intersect with hotspots | RF peak intersection / RF peak length |
| --- | --- | --- | --- | --- | --- |
| S | CD8+ | HLA-I | 0.412<br>($p = 2.51e-53$) | 0.842 | 0.6222 |
| | CD8+ | pan-HLA | 0.236<br>($p = 1.46e-17$) | 0.790 | 0.5497 |
| | CD4+ | HLA-II | 0.201<br>($p = 4.21e-13$) | 0.857 | 0.5305 |
| | CD4+ | pan-HLA | 0.302<br>($p = 2.65e-28$) | 1.000 | 0.6180 |
| N | CD8+ | HLA-I | 0.522<br>($p = 1.46e-30$) | 0.600 | 0.4433 |
| | CD8+ | pan-HLA | 0.507<br>($p = 1.14e-28$) | 0.600 | 0.5957 |
| | CD4+ | HLA-II | 0.560<br>($p = 6.27e-36$) | 0.667 | 0.6463 |
| | CD4+ | pan-HLA | 0.634<br>($p = 2.51e-48$) | 1.000 | 0.5528 |

**Table 2.** *Counts of immunogenic pMHC complexes with peptides originating from key SARS-CoV-2 proteins across 18 COVID-19 patients* (15). From among all pMHC candidates evaluated, several were recognized by T cells across multiple patients. For all proteins except ORF3a, the majority of immunogenic peptides were from locations identified as pan-HLA hotspots.

| Protein | Unique T-cell recognized pMHCs at pan-HLA hotspots | Unique T-cell recognized pMHCs | T-cell recognized pMHCs (across all patients) | pMHC candidates |
| --- | --- | --- | --- | --- |
| S | 14 | 18 | 18 | 616 |
| N | 4 | 4 | 7 | 177 |
| ORF1ab | 60 | 79 | 93 | 1868 |
| ORF3a | 1 | 11 | 13 | 140 |
| M | 5 | 6 | 6 | 138 |
| E | 1 | 1 | 1 | 28 |

**Table 3.**
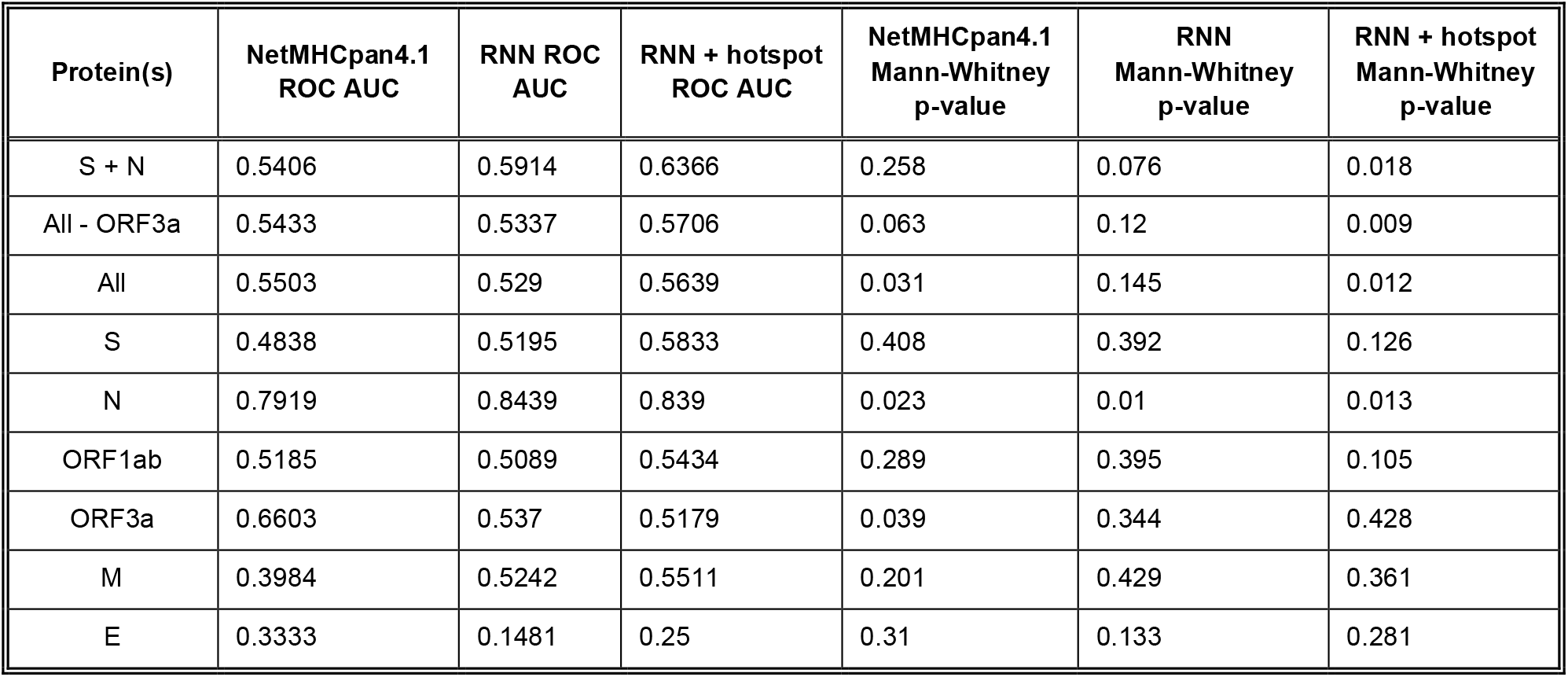
ROC AUC and Mann-Whitney p-values to capture the separability of immunogenic vs. non-immunogenic pMHCs based on NetMHCpan-4.1 rank scores (15) and our RNN binding predictions.

### Verified T-cell epitope response frequencies correlate with aggregated predictions

Grifoni et al. (9) collected confirmed CD8+ and CD4+ epitopes across 25 studies and used the Immunome Browser (26, 27) hosted by IEDB [https://www.iedb.org/] (28) to compute position-specific T-cell response frequency (RF) along with confidence interval (CI) for key viral proteins. Using their provided data, we replicated the RF calculation to validate alignment with our aggregate HLA binding predictions and epitope hotspot locations for both the S and N proteins (**Fig. 2A-F**).

In N, the CD8+ RF lower bound (CI at 95%) correlated with our aggregated HLA-I predictions with Spearman *r_s_* = 0.522 (p = 1.46e-30); and with our pan-HLA aggregate with *r_s_* = 0.507 (p = 1.14e-28). The correlation of the CD4+ RF lower bound in N was at *r_s_* = 0.560 (p = 6.27e-36) and *r_s_* = 0.634 (p = 2.51e-48) with our HLA-II and pan-HLA predictions, respectively. For the S protein, the CD8+ RF lower bound correlates with our HLA-I and pan-HLA aggregated predictions at *r_s_* = 0.412 (p = 2.51e-53) and *r_s_* = 0.236 (p = 1.46e-17); and the CD4+ RF lower bound correlates with our HLA-II and pan-HLA predictions at *r_s_* = 0.201 (p = 4.21e-13) and *r_s_* = 0.302 (p = 2.65e-28).

Note that the lower bound CI for RF is a function of number of subjects for which a certain protein position was interrogated. Thus we do not expect perfect correlation due to non-uniform coverage across studies, as well as the understanding that HLA binding is necessary but not sufficient for T-cell response. Further, the HLA composition of aggregated studies (9) is limited by the regions in which each study was performed, and thus not necessarily reflective of the same global distribution used to inform our HLA set. However, these correlations indicate that when binding predictions are aggregated across a diverse set of HLAs and peptide lengths, the resulting representation of binding promiscuity is useful for predicting immunodominant protein regions yielding peak T-cell epitope frequencies across large samples of subjects.

Although correlation of aggregated HLA predictions with RF lower bound was lower in S than N, significant peaks of the RF were effectively captured in both proteins even when relative magnitudes differed. Employing the same methodology used to select our potential epitope hotspots on the unsmoothed RF lower bound, we found that the majority of regions identified based on the RF signal were indeed covered by our predicted epitope hotspot regions (**Table 1**).

To layer insight from empirical epitope evidence across studies on top of our predictive analysis, we ranked our predicted epitope hotspots according to the maximum RF lower bound within each hotspot range (**Fig. 3**). This helped interpret our subsequent results with an observation driven approach to prioritizing the impact of HLA binding changes that lead to potential for epitope loss.

**Figure 3.**
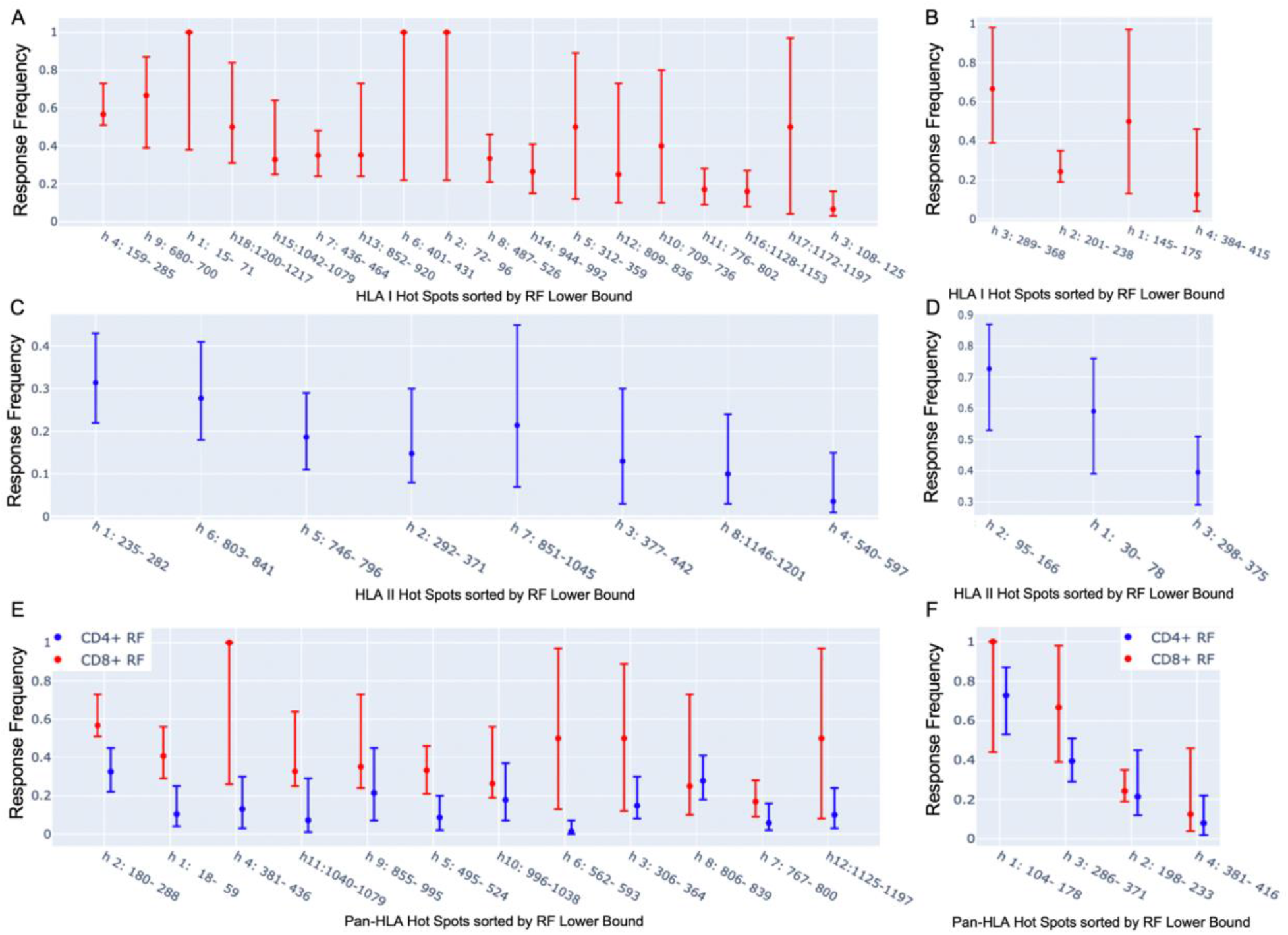
Observation-driven prioritization of epitope hotspots. Shown are our potential epitope hotspots based on binding predictions aggregated across HLAs and peptide lengths, sorted according to the maximum position-specific response frequency (RF) lower bound (9) within their ranges. We ranked HLA-I hotspots according to CD8+ RF (A and B), HLA-II hotspots according to CD4+ RF (C and D), and pan-HLA hotspots according to CD8+ RF (E and F), for S and N, respectively, for each. Each plot illustrates the maximum RF within hotspot ranges, as well as the max of both the 95% confidence interval upper and lower bounds.

### Verified CD8+ T-cell epitopes coincide with predicted epitope hotspots

We also validated predictions against an individual study in which DNA-barcoded peptide-MHC complex (pMHC) multimers were used to identify 122 unique epitopes recognized by SARS-CoV-2-specific CD8+ T cells across 10 HLA-I molecules in 18 COVID-19 patients (15). Of the total unique epitopes from across the viral genome, 118 unique epitopes (119 unique pMHC complexes) were found to originate from the set of 6 proteins considered in **Figure 1**. The most frequent sources of epitopes reported were the ORF1, S, and ORF3 proteins. Further, 4 immunodominant epitopes (recognized by T cells in > 50% of analyzed patients) were found to come from ORF1 and one from N.

For all proteins illustrated in **Figure 1** except ORF3a, the majority of verified epitopes occurred at our pan-HLA potential epitope hotspots (**Table 2**). As previously shown with NetMHCpan-4.1 rank score results (15), we also verified that immunogenic pMHC complexes were dominated by high-confidence binding predictions from our system (**Table 3**). Following the prior example of Saini *et al*. (15), we reported Mann-Whitney test p-values when comparing predictions for immunogenic pMHCs versus those that did not trigger immune response. Due to the equivalence of the Mann-Whitney test with ROC AUC, the latter was also included for clarity. Note that both our classifiers and NetMHCpan-4.1 were trained to predict MHC binding, whereas the ROC AUC scores in **Table 3** reflect how much predicted binding contributes to separability of immunogenic vs. non-immunogenic peptides within the 18 patient samples in the study.

If hotspot selection had no benefit for the identification of immunogenic peptides, then restricting the set of non-immunogenic examples to only those outside of pan-HLA hotspots when computing ROC AUC (or the Mann-Whitney test) would not impact the score. However, we found that in these scenarios, ROC AUC consistently increased (**Table 3**); indicating that hotspot selection is in fact a helpful heuristic for rejecting peptides likely to be non-immunogenic.

Notably, when considering the S and N proteins on which we focused our surveillance, the impact of pan-HLA hotspot selection enhanced separability from an ROC AUC of 0.5914 (when all RNN predictions were considered), to 0.6366 after negatives were restricted to samples outside of hotspot locations. As a baseline comparison on these two proteins: NetMHCpan-4.1 rank values (with no hotspot selection) led to an ROC AUC of 0.5406.

Since MHC binding does not guarantee immunogenicity, it is not surprising that at individual peptide resolution these scores were low compared to what one might expect when a classifier is trained and tested on the same task. It is precisely because individual peptide-MHC binding predictions are not directly predictive of immunogenicity - but are a prerequisite - that we focus on a holistic perspective of binding loss at critical locations as an approach to track and highlight viral variants with most potential for T-cell epitope loss.

### Mutations in S and N protein across SARS-CoV-2 genomes frequently overlap potential epitope hotspots

At bi-monthly time points throughout 2021, a comprehensive analysis was performed of HLA-I and HLA-II binding predictions for S and N proteins across all available SARS-CoV-2 genomes provided by NIH NCBI. In order to focus on impactful variations and minimize erroneous samples, only proteins whose amino acid sequences were observed at least 3 times were considered.

Potential epitope hotspots were mapped from the reference protein sequences to each unique version of S and N to enable direct comparison of predicted binder counts at hotspots. **Figure 4** summarizes the fraction of unique versions of the S and N proteins that are impacted by hotspot mutations from the reference strain, NCBI Reference Sequence NC_045512 (23). This alignment led to the observation that a majority of unique versions of both S and N proteins included at least one mutation at a T-cell epitope hotspot region. For example, in data from July 27, there were 3752 unique versions of the S protein, of which a fraction of 0.868 had mutations at HLA-I hotspots, 0.686 had mutations at HLA-II hotspots, and 0.779 had mutations at pan-HLA hotspots. At the earlier time point of March 19^th^, there were only 1081 unique versions of S and a fraction of 0.691 had HLA-I hotspot mutations, 0.398 had HLA-II hotspot mutations, and 0.563 had pan-HLA hotspot mutations. In the first half of 2021, as the pandemic progressed, the fraction of unique proteins that included mutations impacting hotspots continued to increase rapidly. It finally plateaued in the latter half of 2021, at a state where an overwhelming majority of unique proteins included mutations at epitope hotspots. This suggests that evolutionary pressure favored viral variants with mutations in S and N proteins at regions we identified as HLA binding hotspots. We caution that the causal mechanism for such a pressure is likely to be linked to a multitude of factors outside of T-cell recognition of epitopes alone.

**Figure 4.**
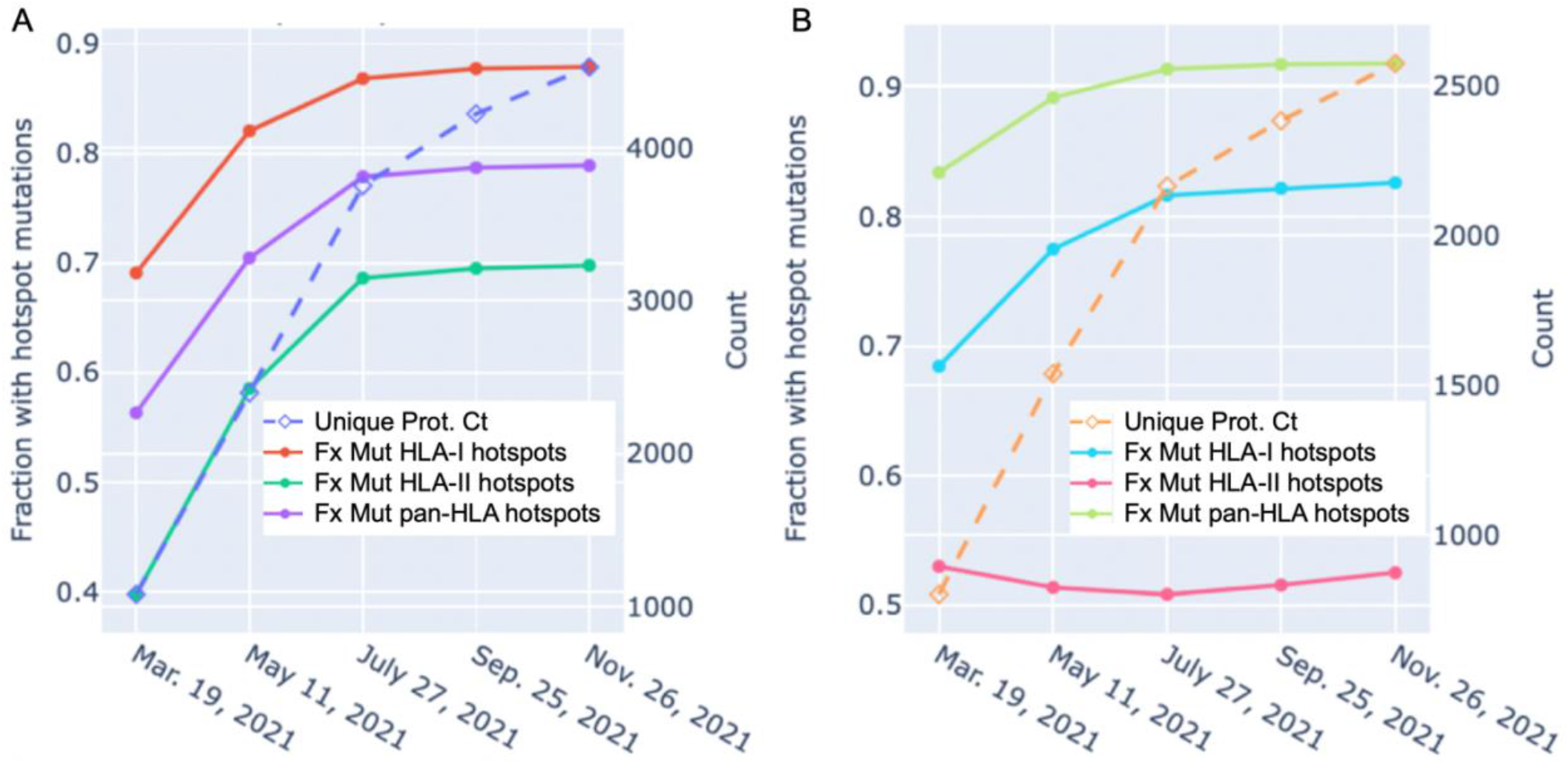
Unique protein count and fraction of unique proteins with hotspot mutations over time for spike (S) and nucleocapsid (N) proteins. The unique protein count (right y-axis) and fraction of unique proteins with mutations at HLA-I, HLA-II, and pan-HLA hotspots (left y-axis) as shown for S (A) and N (B). Unique protein counts included all versions of a viral protein whose amino acid sequence appeared at least 3 times in the viral genomes available through NIH NCBI at bi-monthly time points throughout 2021. Every unique protein that included at least one mutation relative to the SARS-CoV-2 reference genome occurring within an HLA binding hotspot was counted in the fraction of unique proteins with hotspot mutations.

A noteworthy counter example to the above trend are versions of the N protein with mutations at HLA-II hotspots; the fraction of which has stayed steady throughout 2021. The key distinction between HLA-II hotspots identified in N is that regions in the middle and end of the protein that were prominently predicted as promiscuous HLA-I binders (and included in HLA-I and pan-HLA hotspots) were predicted to have minor success binding with HLA-II, and thus did not pass hotspot selection thresholds for class II (**Fig. 1D, E, and F**). The minor contribution of T-cell epitopes from regions omitted in our HLA-II hotspots was confirmed by empirical CD4+ epitope response frequency data (**Fig. 3D and F**). Overall, we did not observe the emergence of mutations in the N protein over the course of 2021 that significantly affected predicted N-directed CD4+ T-cell epitope repertoires.

### Sorting protein variants by fraction of binders lost across representative HLAs

To track the evolution of predicted binding loss - thus potential for epitope loss - across viral variants, we counted the number of predicted binders at each pan-HLA hotspot for every unique version of S and N. This enabled us to measure the fractional change in binding between any two versions of a protein at any specific hotspot location, or in aggregate over all hotspots.

Interactive visualizations illustrating per-HLA aggregate binder count fraction relative to the reference SARS-CoV-2 genome (NCBI Reference Sequence NC_045512) for all S and N epitopes have been made available at https://research.immunitybio.com/scov2_epitopes/, and a static example is illustrated in **Figure 5**.

**Figure 5.**
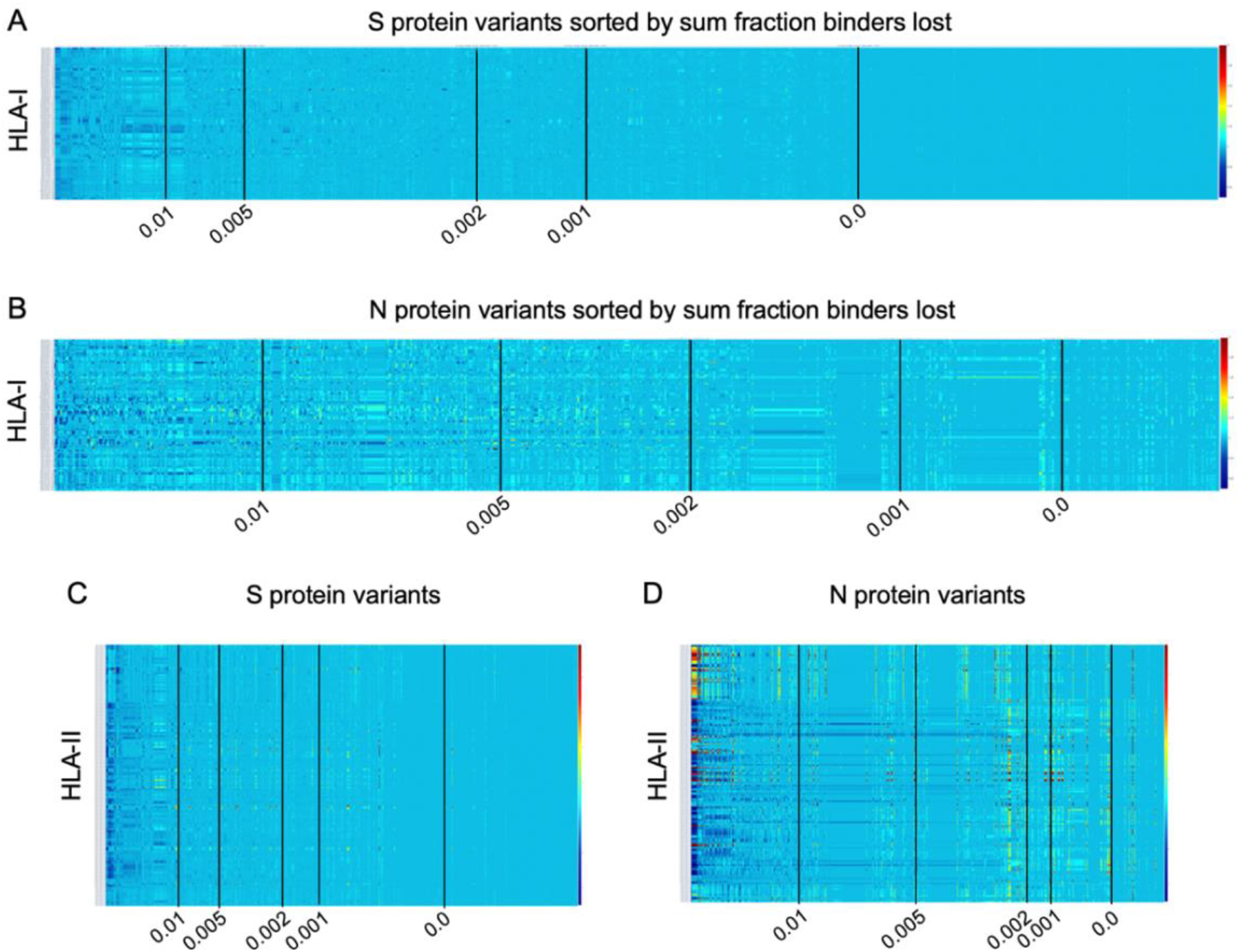
Illustration of per-HLA aggregate binder count fraction at pan-HLA hotspots illustrated for all HLAs across all unique versions of S and N. All plots were sorted according to *sum fraction of binders lost*, such that proteins with the most loss appear on the left. Each plot has 5 vertical black lines indicating average bind loss per HLA thresholds at 0.01, 0.005, 0.002, 0.001, and 0. For compactness, binder count fractions in data only from March 13, 2021 are shown here: HLA-I binder count fractions for all unique versions of (A) S and (B) N; and HLA-II binder count fractions for (C) S and (D) N variants. Interactive plots from all data sample times are available online.

Independently for our comprehensive HLA-I and HLA-II sets, all versions of the S and N proteins were ranked according to *sum fraction of binders lost* (see *Methods* and **Fig. 5**) such that protein versions with the most significant overall loss in HLA binders relative to the reference genome were ranked above protein with less epitope loss potential. A fractional value (*bind loss rank fraction*) was assigned to represent ranking position; where values near 0 correspond to the top of the ranking order (most loss of HLA binders), and values near 1 indicate a rank at the end (least loss of HLA binders). This was done at each bi-monthly data time point. Since the number of unique proteins increased through time and therefore impacted relative rankings, to enable comparison of lineages between sample times *sum fraction of binders lost* was divided by the number of HLAs considered to obtain an *average bind loss per HLA*.

To prioritize browsing protein versions with most potential for epitope loss, all interactive plots were sorted as described above. Visualizations were also supplemented with *average bind loss per HLA* thresholds drawn at 0.01, 0.005, 0.002, 0.001, and 0 to clearly delineate zones where binding loss is likely to have little or no impact.

### High fraction of lineages with the most binder loss in S show conserved HLA binding in N

Motivated by vaccine approaches currently in trials that include both the S and N proteins (20–22), we investigated the relationship between HLA binder loss in S and N across all viral lineages in the data.

Note that one viral lineage is often defined to span multiple alternative versions of each protein. Conversely, one specific protein sequence may be produced by several viral lineages. To summarize results, each lineage was represented by the sample with worst case potential epitope losses in S and N proteins (highest *sum fraction of binders lost*), and alternatively by the most frequent instance of each protein occurring across genomes assigned to a single lineage classification. Worst case scenarios were considered independently so that *bind loss rank fraction* values assigned to a lineage were allowed to come from different genomes for S and N. This assumed that versions of the lineage may already exist outside the data, or come to exist through continued evolution, where proteins with most significant loss co-occur in the same viral genome. Most frequent versions of S and N representing each lineage were also selected independently.

**Figure 6** shows the relationship between lineages with representative versions of the S protein ranked at the top according to most significant loss in HLA-I or HLA-II binders (*bind loss rank fraction*) and different thresholds of HLA binder loss in N (*average bind loss per HLA*). In data analyzed from November 26, it was found that lineages whose worst-case S protein ranked within the top 1% most potential for epitope loss, 17.5% had an *average bind loss per HLA* less than 0.005 in N for HLA-I (**Fig. 6A**). Further, as the scope was increased to consider lineages with S ranked in the top 10% most epitope loss potential, 33.3% of those lineages demonstrated the same level of conservation in N (**Fig. 6A**). The picture was similarly optimistic when it came to conservation of potential CD4+ epitopes. The same level of HLA-II binder conservation (*average bind loss per HLA* < 0.005) in N was exhibited by 26.5% of lineages that fell in the top 10% according to binder loss in S (**Fig. 6C**). When a stricter requirement was imposed that an equal threshold of binder conservation be observed across both HLA-I and HLA-II in N, there was little change in outcome: the strict N conservation criteria was satisfied by 19.1% of lineages in the top 10% of those with most loss in S (**Fig. 6E**).

**Figure 6.**
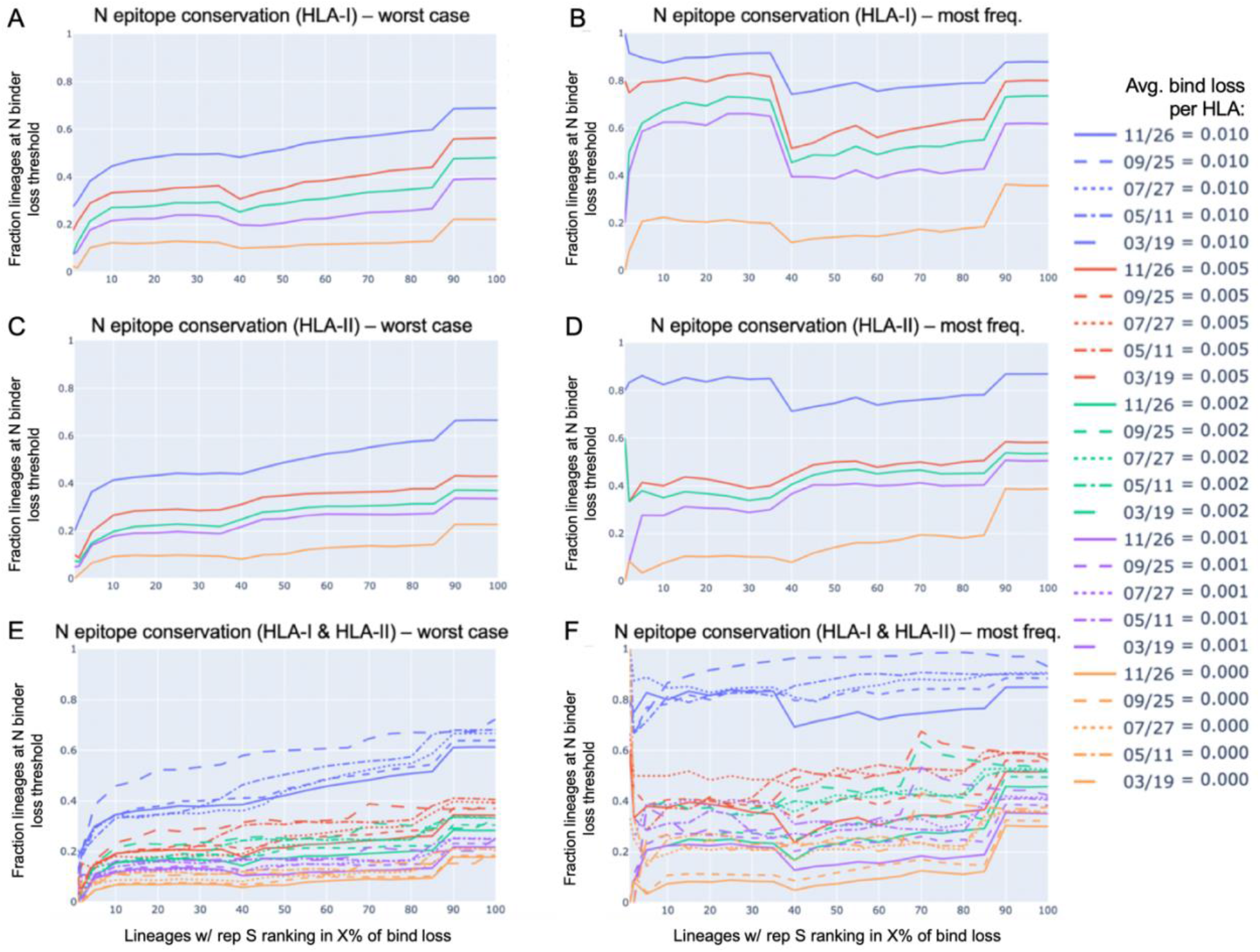
N epitope conservation in SARS-CoV-2 lineages with representative S protein ranking in the top percent potential epitope loss. All viral lineages within the data were represented either by the S and N protein exhibiting the worst-case epitope loss potential (A, C, and E), or by the most frequently occurring versions of S and N (B, D, and F). For lineages with the S protein ranking in the top percent for the most significant loss in HLA-I or HLA-II binders (x-axis), shown are the fractions of those top loss lineages that exhibited (A,B) HLA-I binder conservation in the N protein, (C, D) HLA-II binder conservation in the N protein, or (E,F) both HLA-I and HLA-II binder conservation in the N protein; all at various conservation thresholds of average (avg.) bind loss per HLA, as represented by the colors shown in the legend. The strictest N conservation criteria was used to illustrate the dynamics of the relationship throughout the span of the 2021 (E,F).

In addition to being robust to a strict requirement of binder conservation across both HLA-I and HLA-II, this conclusion also held robustly throughout the course of the pandemic in 2021. Looking at the lineages with worst case S demonstrating binder loss within the top 10% at data sample times in March, May, July, September, and November; the percentage of lineages satisfying the same N conservation criteria as above was 23.5%, 19.3%, 18.1%, 18.8%, 19.1%, respectively (**Fig. 6E**).

Even more compelling is the observation that the most commonly occurring versions of S and N within each lineage (**Fig. 6B, D, and F**) were not typically the versions exhibiting the worst-case HLA binder loss (**Fig. 6A, C, and E**). Lineages with their most frequent S ranking within the top 10% for binder loss, demonstrated strict HLA-I and HLA-II binder conservation in N (at *average bind loss per HLA* < 0.005) at percentages of 40.0%, 37.5%, 50.0%, 40.5%, 37.5% across time (March, May, July, September, and November respectively) (**Fig. 6F**).

### Variants of Concern (VOC) and Variants of Interest (VOI) over time

Our global analysis was leveraged to highlight the evolution of the peptidome in VOC as defined by the Centers for Disease Control and Prevention (CDC): B.1.17 (World Health Organization (WHO): Alpha), B.1.351 (WHO: Beta), B.1.617.2 (WHO: Delta), P.1 (WHO: Gamma), and B.1.1.529 (WHO: Omicron). The evolution of VOI was also tracked: B.1.427 and B.1.429 (WHO: Epsilon), B.1.525 (WHO: Eta), B.1.526 (WHO: Iota), and B.1.617.1 (WHO: Kappa). Note that some WHO labels such as Beta or Delta can correspond to multiple PANGO lineages, thus the complete history of Delta may not be fully captured only by its originating PANGO lineage B.1.617.2.

To supplement NIH NCBI data from November 26, 2021 to cover the emergence and rapid spread of the B.1.1.529 (Omicron) lineage in late 2021, we obtained all Omicron genomes from GISAID (29–31) up to December 6, 2021. All unique versions of S and N proteins that included less than 1 percent unspecified amino acids and were classified by GISAID as Omicron were appended to the November 26 data. This augmented dataset is labeled as December 6 throughout figures. We also later verified that according to all GISAID data available up to two additional time points (December 20, 2021 and December 29, 2021), the versions of Omicron S and N proteins identified as the top two most frequent remained unchanged in their rankings.

With each of the VOC and VOI lineages, we observed that as both overall sample count and variety of unique proteins increased over time for S and N (**Fig. 7A and B; and Fig. 8A and B**, respectively), there was a distinct correlation with increased worst-case HLA-I and HLA-II binder loss (**Fig. 7C-F and Fig. 8C-F**, respectively). As a result of this evolution, lineages increased their chances of successful CD8+ and CD4+ T-cell evasion. However, we also observed that this did not occur at the same rate in all lineages. For example, worst case HLA-I binder loss in S changed extremely rapidly between July 27 and September 25 for both VOC B.1.617.2 (Delta) (**Fig. 7C**) and VOI B.1.429 (Epsilon) (**Fig. 8C**) compared to any other variants. We also saw from the emergence of B.1.1.529 (Omicron) that novel strains can feature high *average bind loss per HLA* in the context of existing lineages. The most frequent versions of B.1.1.529 proteins exhibited the highest potential for epitope loss compared to the most frequent proteins corresponding to almost all other VOCs (**Fig. 7C, D, and E**). The one exception was a close match in *average bind loss per HLA-II* with B.1.1.7 in the N protein (**Fig. 7F**). Emergence of novel variants and differences in rate of evolution occasionally impact the relative rankings of linages according to most epitope loss potential, and thus underscore the importance of continued genomic surveillance.

**Figure 7.**
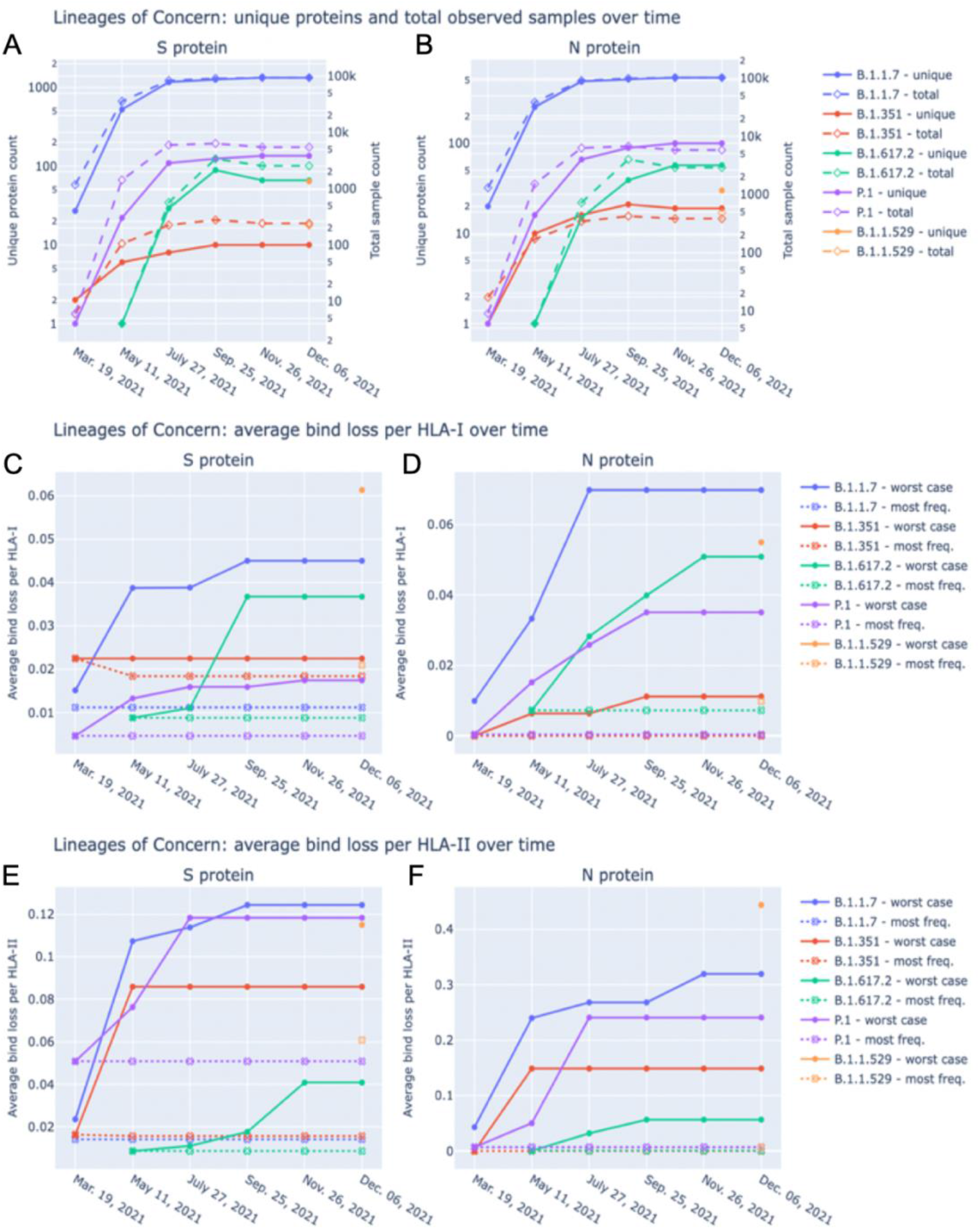
VOC lineage sample counts, unique proteins, and average bind loss per HLA-I and HLA-II across time. Total sample count (right y-axis) as well as the number of unique versions (left y-axis) of S (A) and N (B) proteins classified as VOC lineages are shown at each data sample time. December 6 included November 26 data plus supplementary genomes from GISAID to capture the emergence of B.1.1.529 (WHO: Omicron). Average bind loss per HLA-I (C, D) and HLA-II (E, F) were tracked across time for both scenarios where lineages were represented by their worst-case bind loss versions of S and N (solid lines in C-F), and most frequently occurring versions of S and N (dotted lines labeled most freq. in C-F). Legends identify VOC lineages: B.1.1.7 (Alpha), B.1.351 (Beta), B.1.617.2 (Delta), P.1 (Gamma), and B.1.1.529 (Omicron).

**Figure 8.**
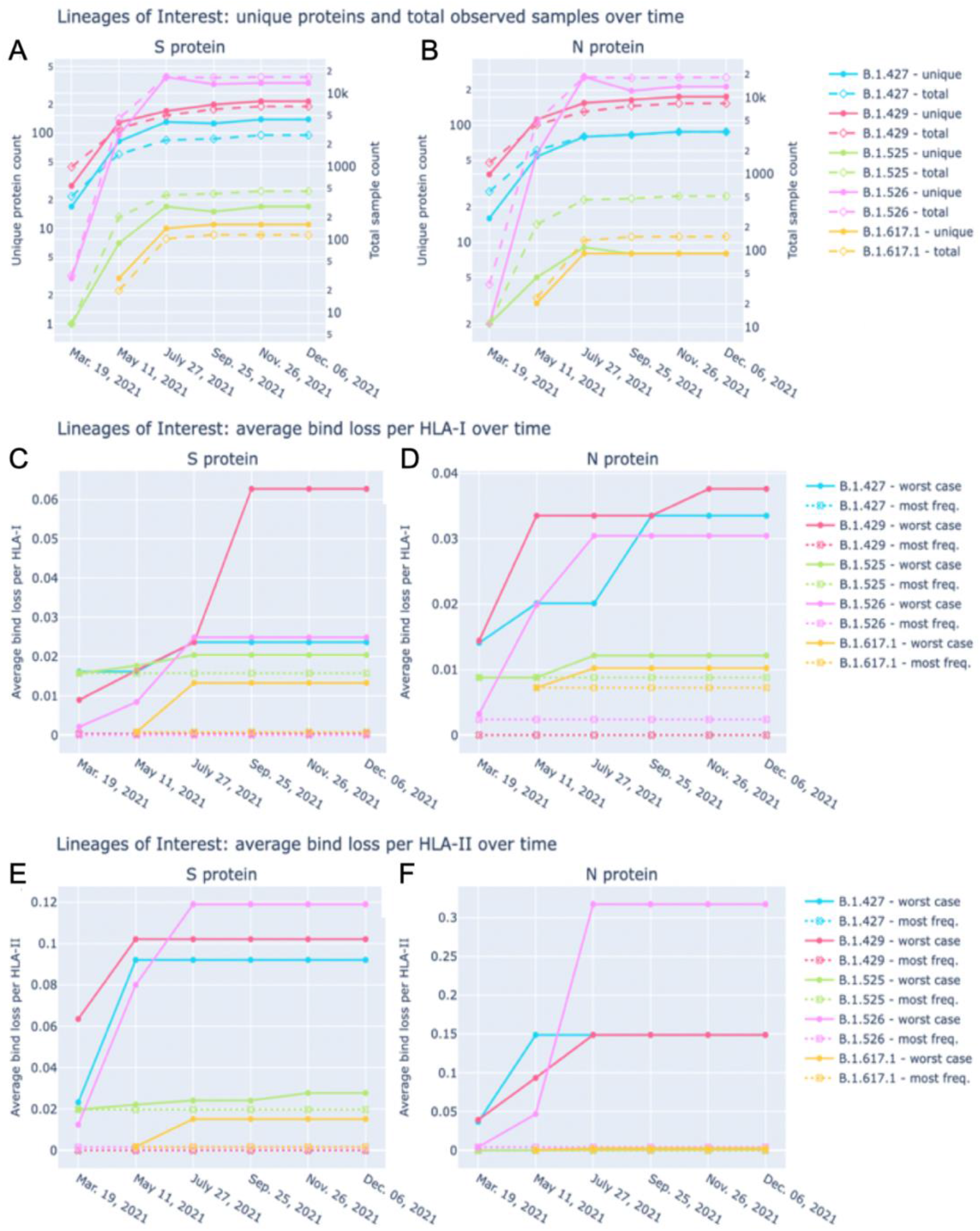
VOI lineage sample counts, unique proteins, and average bind loss per HLA-I and HLA-II across time. Total sample count (right y-axis) as well as the number of unique versions (left y-axis) of S (A) and N (B) proteins classified as VOI lineages are shown at each data sample time. Average bind loss per HLA-I (C, D) and HLA-II (E, F) were tracked across time for both scenarios where lineages were represented by their worst-case bind loss versions of S and N (solid lines in C-F), and most frequently occurring versions of S and N (dotted lines labeled most freq. in C-F). Legends identify VOI lineages: B.1.427 and B.1.429 (Epsilon), B.1.525 (Eta), B.1.526 (Iota), and B.1.617.1 (Kappa).

### VOCs demonstrate increased potential for loss of epitopes when compared to VOI

Across both the S and N proteins, we observed that VOC lineages demonstrate a consistently higher *average bind loss per HLA* than lineages classified as VOI; especially when focusing on the most frequent proteins within each lineage. This held true for VOC versus VOI across both HLA-I (**Fig. 7C and D versus Fig.8C and D**, respectively) as well as HLA-II (**Fig. 7E and F versus Fig. 8E and F,** respectively). In summary, lineages which have spread most successfully, and have been identified as the most concerning, consistently demonstrate a higher rate of epitope loss relative to VOI and other lineages.

### Proteins instances with worst case binding loss did not become the most frequent for any VOC or VOI lineage

Perhaps the most notable observation from our tracking of VOC and VOI lineages is that in no instance did the S or N proteins exhibiting the worst-case HLA-I or HLA-II binding loss become the most frequent protein version for a lineage (**Fig. 7C-F and Fig. 8C-F**). Even though we found an increasing occurrence of mutations at T-cell epitope hotspots over time (**Fig. 4**), genomes with the most potential for T-cell evasion within the trajectory of each lineage were not the most successful at spreading across the population. This appears consistent with hypotheses expressed earlier (7, 10, 13–15), suggesting that T-cell evasion is unlikely to be a primary evolutionary pressure on SARS-CoV-2. Instead of a gradual evolution, we observed step changes in T-cell evasion potential when tracking most frequent versions of S and N within VOCs, as demonstrated by the emergence of B.1.617.2 (Delta) and B.1.1.529 (Omicron) during our sampling period (**Fig. 7C - F**).

### High levels of epitope conservation hold across most frequent versions of N protein in VOC and VOI lineages

For both VOC and VOI lineages, regardless of epitope loss potential in versions of their S protein, a high degree of epitope conservation was observed across nearly all most frequent versions of N. Even lineages whose most common N protein exceeded our earlier defined conservation threshold (*average bind loss per HLA* < 0.005, **Fig. 6**) did so only by a narrow margin: B.1.617.2 and B.1.1.529 (**Fig. 7D**), B.1.617.1 and B.1.525 (**Fig. 8D**) with respective *average bind loss per HLA-I* values: 0.0072, 0.0098, 0.0072, 0.0088; B.1.1.7 and B.1.1.529 (**Fig. 7F**) with *average bind loss per HLA-II* values: 0.0072, 0.0070.

### Localization of significant epitope change across VOC and VOI lineages highlights regions of N conservation

On the S protein the hotspot locations that exhibited some of the most significant potential drops in epitope count across sequenced genomes of VOCs and VOIs also corresponded to regions confirmed to have the highest frequency of T-cell epitopes across aggregated empirical studies (9). Specifically pan-HLA hotspots 2 and 4, which ranked first and third in terms of confirmed epitope response frequency (**Fig. 3E**), were regions where both B.1.351 (Beta) and B.1.1.529 (Omicron) demonstrated notably lower HLA-I binder count fraction values (0.953 and 0.961 at hotspot 2, and 0.922 and 0.938 at hotspot 4; respectively for the two lineages) relative to other VOCs and VOIs (**Fig. 9A**). B.1.617.2 (Delta) also stood out at pan-HLA hotspot 5 with a low HLA-I binder count fraction of 0.958 relative to others. However, hotspot 5 ranked in the middle according to empirical response frequencies.

**Figure 9.**
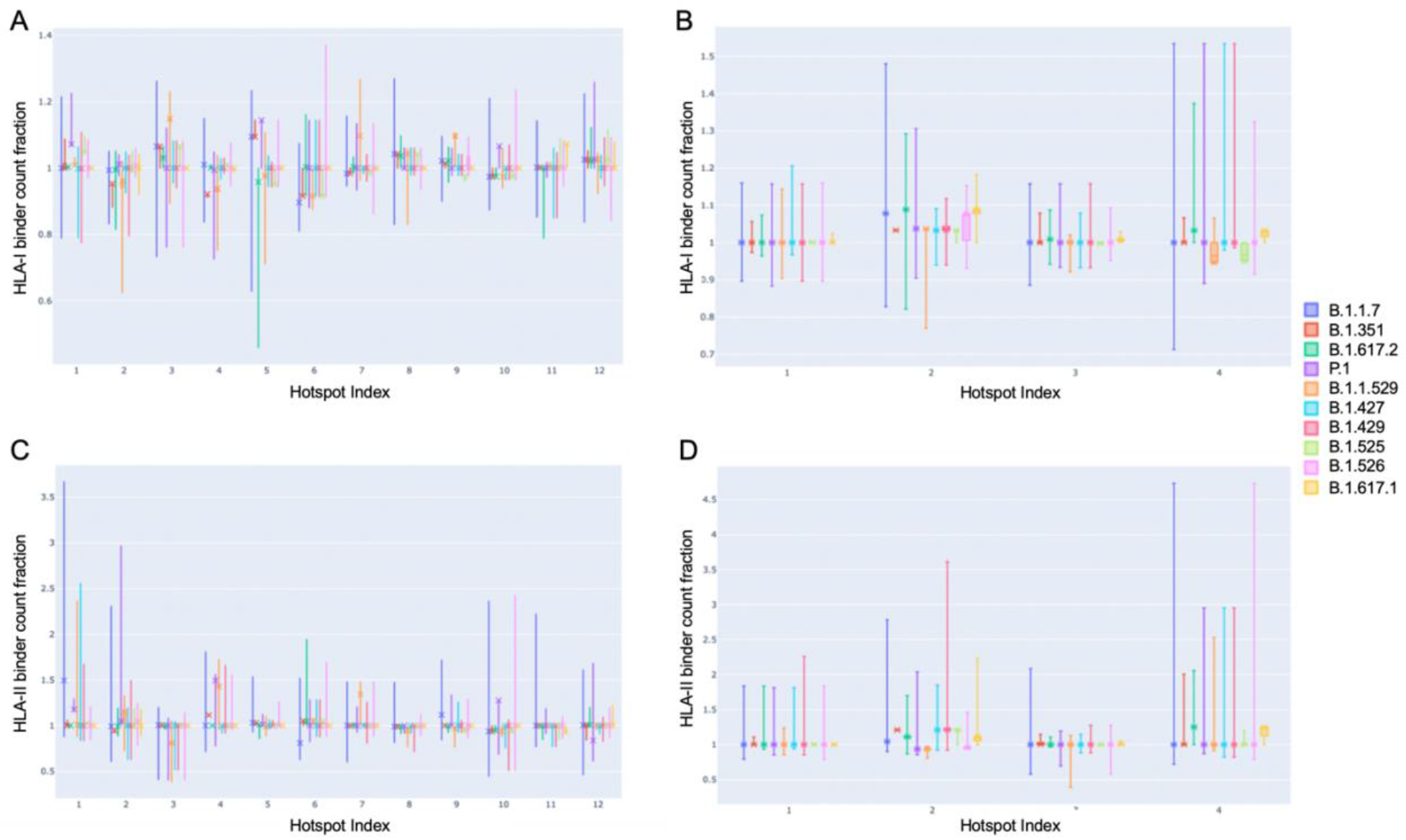
Distribution of binder count fraction relative to reference SARS-CoV-2 at pan-HLA hotspots for VOC and VOI lineages. The box plots illustrate the distribution of HLA-I (A,B) and HLA-II (C, D) binder count fraction averaged across HLAs (y-axis) at S (A, C) and N (B, D) pan-HLA hotspots (x-axis) for all unique proteins classified as VOC and VOI lineages. Binder count fractions specific to the most frequent protein versions of each lineage are annotated with an ‘X’ on top of the box plots. VOC and VOI lineage labels are indicated in the legend at right.

In terms of HLA-II binder count fraction across S protein hotspots, only Omicron stood out at hotspot 3 with a distinctly lower value than other lineages (**Fig. 9C**). Considering hotspot 3 is toward the end of rankings according to response frequency, CD4+ epitopes may, in general, be less subject to T-cell evasion across VOCs and VOIs than CD8+ epitopes.

For N protein, pan-HLA hotspots 1 and 3 were verified as the highest frequency contributors of both CD4+ and CD8+ epitopes according to empirical evidence (**Fig. 3F**). As previously discussed, we were able to infer that these hotspots were conserved, that is, less subject to mutation in the course of viral evolution (**Fig. 4B**); and - as expected - we observe that they are the most stable across all variants in terms of HLA-I and HLA-II binder count fraction at or near 1 (**Fig. 9B and D**).

### VOCs demonstrate heterogeneity of epitope loss potential at pan-HLA hotspots across our comprehensive HLA set

Finally, to illustrate that epitope loss is not uniform across HLAs, we visualized the HLA-I binder count fraction relative to reference proteins at pan-HLA hotspots for the most frequent S (**Fig. 10A-E**) and N (**Fig. 10G-K**) proteins of all VOCs. We also included the second most frequent instances of Omicron S (**Fig. 10F**) and N (**Fig. 10L**). Again the HLA binding stability of N hotspots 1 and 3 was apparent across HLAs, even when not averaged as in **Figure 7B**. In contrast, multiple hotspots on the S protein demonstrated significant diversity in binder count fraction across our HLA-I set (**Fig. 10A-F**). This was consistent with findings that S regions yielding CD8+ epitopes vary with patient repertoire of HLA-I alleles, and thus yield a more heterogeneous distribution of position specific response frequency in S across aggregated studies (14). For closer inspection, interactive visualizations of **Figure 10** are available online along with other supplementary content.

**Figure 10.**
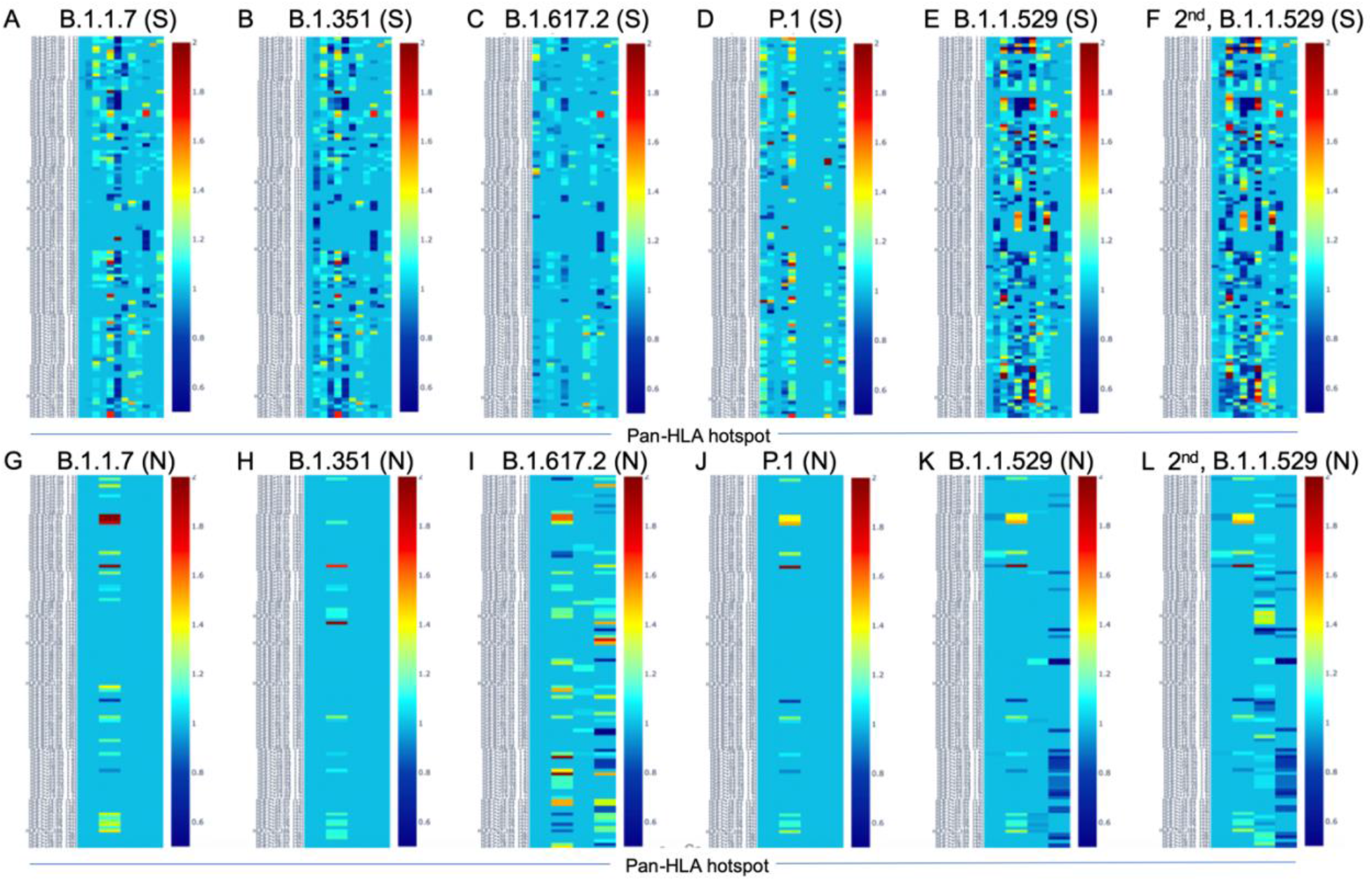
Change in HLA-I binder count from SARS-CoV-2 reference at pan-HLA epitope hotspots for most frequent S and N proteins of VOC lineages. Change in potential epitope count as a fraction of reference count is shown for all HLA-I in our analysis set (y-axis) for the most frequent versions of S (A-F) and N (G-L) in each of the VOC lineages: (A,G) B.1.1.7 (Alpha), (B,H) B.1.351 (Beta), (C,I) B.1.617.2 (Delta), (D,J) P.1 (Gamma), (E,K) B.1.1.529 (Omicron), as well as the second most frequent versions of Omicron proteins (F,L). For both S and N proteins pan-HLA hotspots (x-axis) were used. Interactive versions of all plots above are available online.

## Discussion

With our unique approach to integrating HLA binding predictions across a broad range of peptide lengths and a representative HLA set, we were able to demonstrate that our aggregate signals representing candidate epitope frequency and HLA promiscuity correlated well with empirical evidence of T-cell response frequencies, and correctly identified protein regions yielding immunodominant epitopes across multiple different cohorts (9, 14, 15) (**Figs 1 and 2; Tables 1, 2, and 3**). It is particularly noteworthy that multiple prior studies had omitted predictive analysis when investigating CD4+ T-cell reactivity, attributing this omission to HLA-II binding prediction algorithms not effectively predicting epitope recognition (14). Our results have demonstrated that our binding prediction algorithms coupled with our approach to integrating predictions across lengths and HLAs were similarly effective at identifying regions of both immunodominant CD4+ epitopes and CD8+ epitopes (**Fig. 2; Table 1**).

Our approach to selecting representative HLA sets also benefitted from our binding prediction models. We leveraged learned HLA representations to construct a global HLA set based not only on population frequency, but also on uniqueness of function. The goal was to ensure we did not miss consideration of HLAs that may not be frequently studied or well characterized in datasets, but were predicted to be distinct from other HLA groups in how they interact with peptides.

In contrast to in-silico analyses of singular (9, 13) or select (13, 32) sequences derived from representative viral genomes or impacts of individual mutations, our focus on full protein sequences obtained from all available viral genomes at each sample time enabled us to capture deeper co-occurrence relationships between mutations and their impact on epitope loss potential within and across lineages. This yielded a unique view of viral evolution from the perspective of potential for T-cell evasion.

Notably, we observed that throughout 2021 there were no cases among VOC or VOI lineages where the S or N protein versions with the most epitope loss potential became the most frequently observed within a lineage. This occurred despite trends of increasing rates of mutations at potential epitope hotspots overall (**Fig. 4**), and worst-case protein versions exhibiting more epitope loss within lineages as time progressed (**Figs 7 and 8**). This suggests that T-cell evasion is not a dominant evolutionary pressure on SARS-CoV-2 evolution, in agreement with prior hypotheses (7, 10, 13–15).

When looking across all SARS-CoV-2 lineages, we observed that among lineages with their S protein demonstrating worst case binder loss within the top 10%, 19.1% had a high level of epitope conservation in their worst-case version of N. This relationship proved even more distinct when representing lineages by their most frequent versions of S and N; boosting the fraction of lineages with N conservation to 37.5% among lineages with their S in the top 10% of binder loss. We observed that these relationships remained stable over time throughout the course of the pandemic, even as new variants emerged and existing lineages continued to mutate (**Fig. 6**).

Delving deeper, we found that predicted hotspots aligning with the highest frequencies of empirically verified epitopes were among the most impacted by HLA binding loss on the S protein. Whereas on the N protein, the top two epitope contributing hotspots happened to be conserved across time, showing almost no loss in HLA binding (**Fig. 9**). We further found that candidate epitope regions most impacted by N protein mutations were specific to CD8+ T cells and that the virus did not show an accelerated the rate of mutation that significantly impacted the predicted CD4+ T-cell epitope repertoire of the N protein over the course of 2021.

A key limitation of this study is that our analysis is built upon predictions of peptide binding to HLA-I and HLA-II molecules, which is known to be necessary but not sufficient, for T-cell recognition. Not all predicted binders are likely to be T-cell epitopes, and thus not all changes in HLA binding prediction are guaranteed to lead to epitope loss. We aimed to minimize this disparity with our approach to integrating predictions across peptides, lengths, and HLAs; and by focusing on relative change at hotspots of potential epitopes. But ultimately there are unmodeled factors between HLA presentation and T-cell response, and an additional layer of predictive tools trained directly for the task may be necessary to improve precision in the future.

A secondary limitation to the analyses presented herein is that our conclusions are based around a representative set of HLAs selected to cover the most general picture of the world population (and be inclusive of any minority but distinctly functioning HLA groups). Thus conclusions may vary if the analysis were to be repeated with a different HLA set, for example one specific to a population or region. We also encountered very sparse coverage in population statistics of HLA-II alpha-beta haplotypes. To overcome this, we selected our HLA-II set based on all individual alpha and beta chain frequencies and considered all permutations, and may have incurred a risk of overrepresenting the impact of some alleles.

Finally, it is important to note that our conclusions were based on viral genome data curated by the NIH NCBI and GISAID. Therefore, there may be multitudes of unaccounted factors such as regional variability in available testing, processing times, sequencing methodologies, etc.; which might affect how representative the data was of the true state of the virus at any one point in time.

Overall, our work yielded a novel perspective into the evolving landscape of T-cell evasion potential over the time period studied, and added to the mounting body of evidence suggesting that the inclusion of proteins such as N, along with S, in vaccines is likely to be an effective strategy for improving robustness and duration of immune memory protective against SARS-CoV-2 infection and severe COVID-19 in the face of evolving viral variants.

Interactive visualizations allowing researchers to delve deeper into our data and findings are available at: https://research.immunitybio.com/scov2_epitopes/

## Methods

### HLA-I and -II binding prediction with RNN and CNN

A convolutional neural network (CNN) as well as an attention augmented recurrent neural network (RNN) were developed for HLA-I and HLA-II binding prediction using data from IEDB [https://www.iedb.org/] (28) downloaded in January 2020. Training data included 436,805 unambiguously labeled peptide-HLA examples across 159 unique HLA-I A, B, and C alleles; and 96,818 examples spanning 17 unique HLA-II alpha chains and 73 unique HLA-II beta chains.

Both neural net architectures utilize the full HLA protein sequence as well as the full peptide amino acid sequence as inputs. HLA-I and HLA-II binding prediction systems were trained independently, but weights learned from the best HLA-I systems were used to initialize corresponding architecture components for HLA-II training. Rather than estimating affinity, both systems predict binding as classification. This design choice alleviates issues due to high variance in affinity measurements observed in IEDB data (**Fig. 11**), which required prior systems to add ranking to normalize predictions across HLAs (33–35).

**Figure 11.**
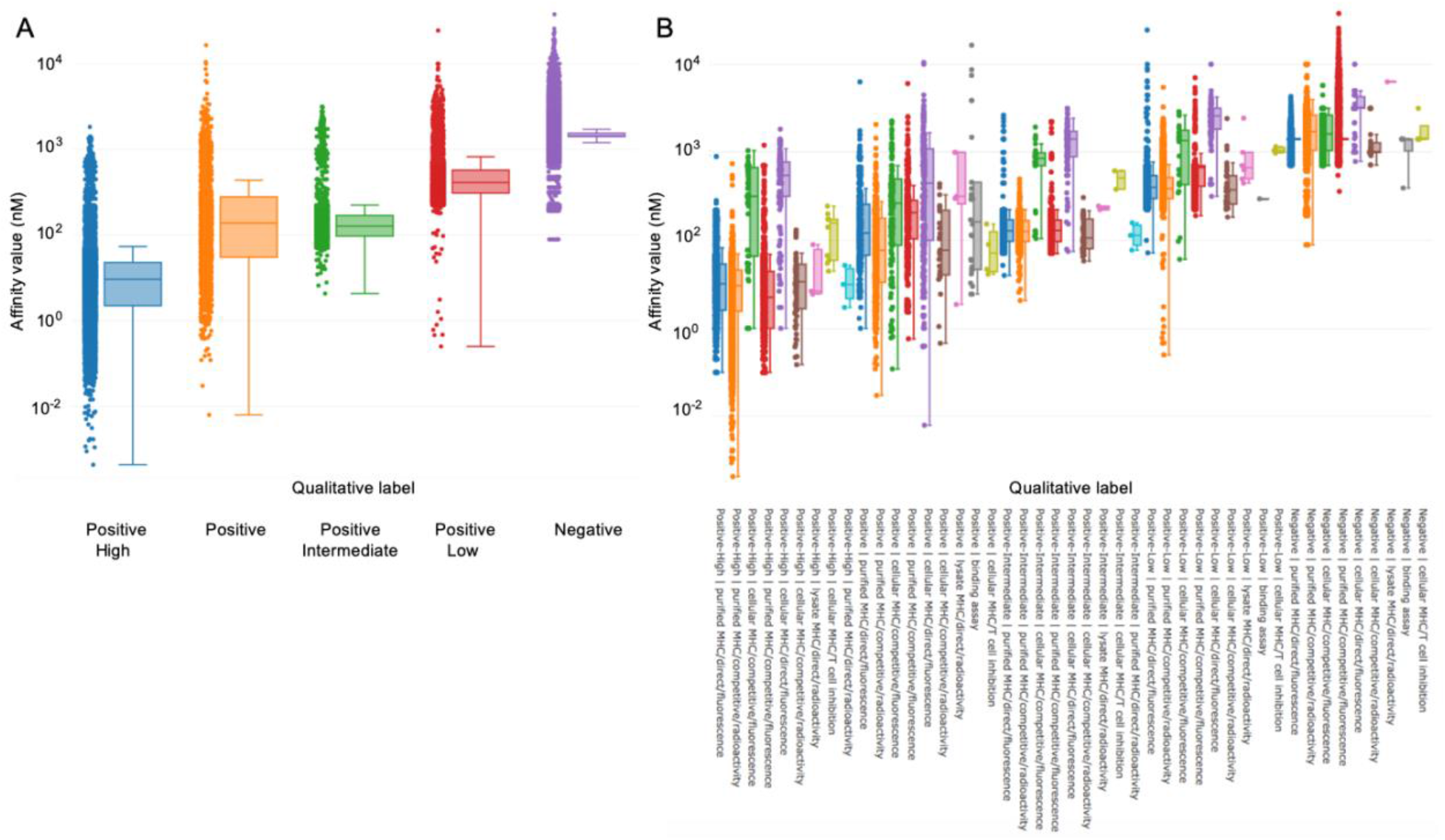
IEDB affinity measurements. (A) IEDB affinity measurements are represented on the y-axis and curated qualitative assessments of binding on the x-axis. (B) Affinity is shown on the y-axis and a qualitative label presented by “assay/method” used to measure affinity on the x-axis. Each assay or method was assigned a distinct color.

As represented in **Figure 11A**, IEDB affinity measurements were found to exhibit high variance across curated qualitative assessments of binding. Although some biases may have been due to measurement method, even when the same assay was used, significant variance in affinity values across binding categories remained (**Fig. 11B**). The commonly selected cutoff of 500 nM for confidently binding peptides is thus likely to mislabel or ignore many valid binders. We therefore expected that classification of peptide binding and presentation was likely to be a better posed problem than attempting to predict extremely noisy binding affinities.

Validating our selection of classification over regression, our RNN classifier’s raw predictions were consistently confident of binding for verified eluted ligands across 27 HLA-I molecules in the Pearson *et al*. dataset (36) (**Fig. 12**). Large variance of predicted affinity values across HLAs in this dataset was the motivation for re-scoring raw NetMHCpan-4.0 predictions (12) by instead reporting HLA specific ranking values. By achieving comparable outcomes to affinity prediction followed by re-ranking, but without needing the additional independent ranking step to re-calibrate predictions, we confirmed that classification based on IEDB’s curated qualitative labels was better posed when considering a multitude of HLAs.

**Figure 12.**
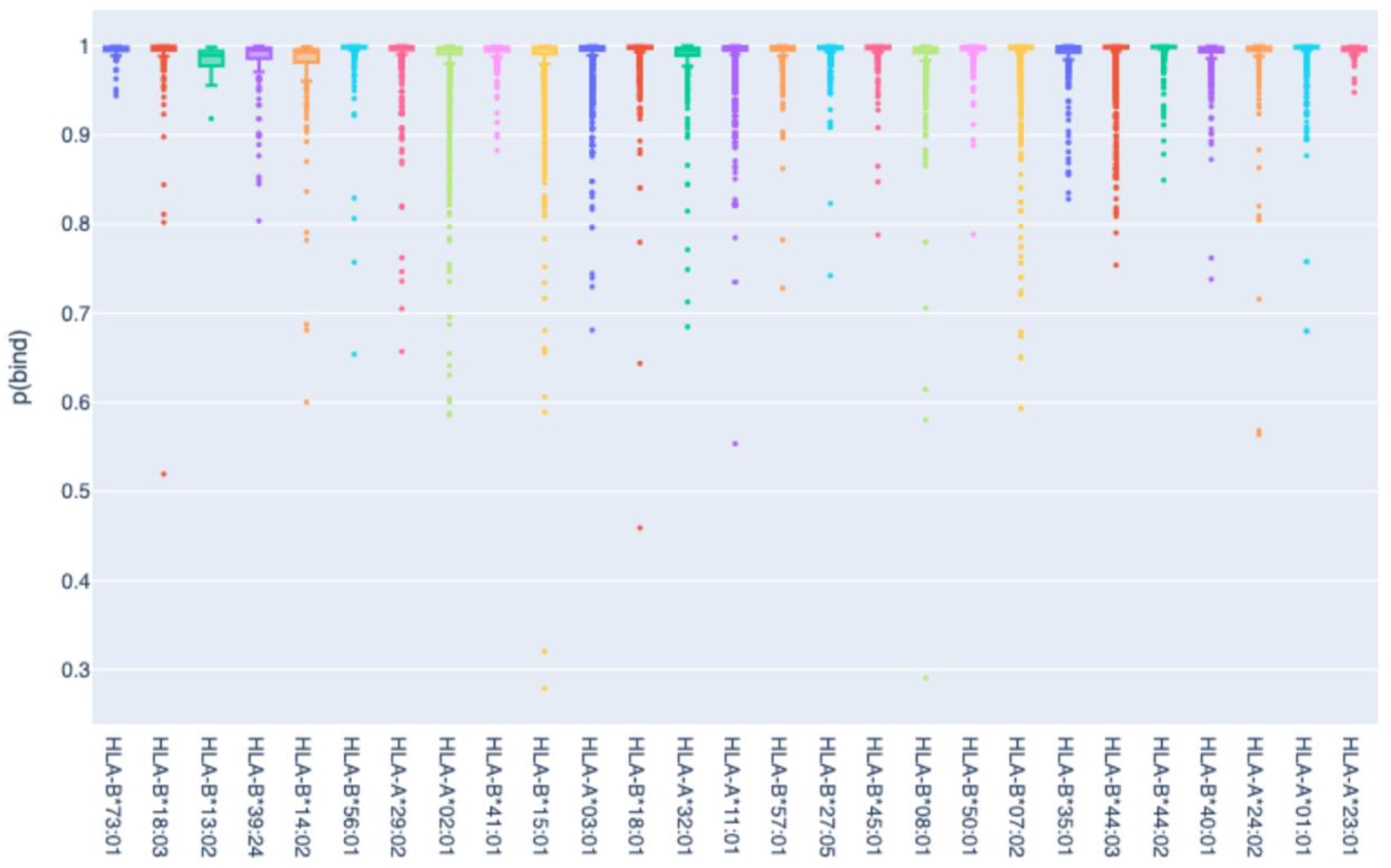
Box plots of all RNN classifier binding predictions across 27 HLA-I molecules. Empirically verified eluted HLA-I ligands in the Pearson *et al*. dataset (36) are shown in the x-axis and our binding prediction values on the y-axis.

During each RNN and CNN training epoch, data for every distinct HLA was augmented with an additional 20 percent background negative peptides (or a minimum of 150 for each HLA-I and 25 for each HLA-II) randomly sampled from the Swiss-Prot human proteome (https://www.uniprot.org/proteomes/UP000005640) and required to not already exist in training.

To help address data imbalance between HLAs, as well as positive/negative sample imbalance within each HLA, we adapted the sample weighting strategy proposed by Cui *et al*. (37) independently across both scenarios (weight smoothing parameter beta = 0.99 was used within each HLA to balance positive/negative examples, and beta = 0.999 was used to balance contributions from each HLA). To ensure average batch loss magnitudes were not impacted by the re-weighting scheme, the combined weights were re-normalized to sum to twice the number of distinct HLAs in training.

Both trained RNN and CNN neural networks were evaluated to compare favorably on the test set of eluted ligands across 36 HLA-I molecules published with the release of NetMHCpan-4.1 (test data and results from other evaluated systems (33, 34, 38, 39) are available at: http://www.cbs.dtu.dk/suppl/immunology/NAR_NetMHCpan_NetMHCIIpan/). To compare directly to the state-of-the-art (34), we used the same methodology to estimate PPV for each HLA: based on the fraction of true positive peptides in the top *N* predictions, where *N* was defined by the total number of true positives for the HLA of interest multiplied by a factor of 0.95. The mean of this PPV estimate across HLAs is reported in Table 4 along with mean ROC AUC.

**Table 4.**
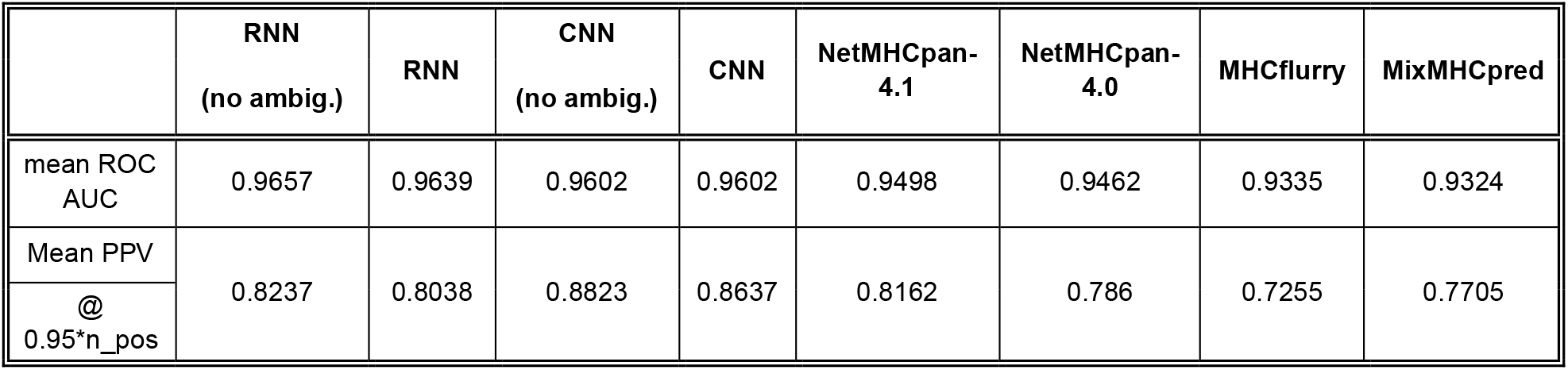
Performance evaluation on eluted ligand test NetMHCpan-4.1, (34)dataset including 77,053 peptides (lengths 8-14) across 36 distinct HLA-I molecules. Our system identifies situations where predictions are ambiguous; columns with the “(no ambig.)” label do not include ambiguous data; this is in contrast to a forced decision in all cases for columns labeled only RNN or CNN.

Our RNN and CNN systems both incorporate an ensemble of 5 models with different random weight initializations and different random training/validation splits, but otherwise identical model architecture and hyperparameters. The variance of predictions from such an ensemble has been shown to be an effective proxy for model uncertainty (40), particularly in cases such as unseen data. Using this additional metric, our models are able to abstain from forcing a classification decision when uncertainty is high. Specifically, ensemble predictions for which the classification decision boundary falls within a standard deviation of the mean are classified as ambiguous and can be disregarded. Additional columns in **Table 4** indicate that this ability to abstain from classification improves accuracy for both the RNN and CNN.

### HLA-I grouping and selection

HLA nomenclature was originated by the *World Health Organization (WHO) Nomenclature Committee for Factors of the HLA System*, which issued its first report in 1968 based on serology analysis (41) and has continued to refine the resolution of its methods by incorporating molecular and genetic information (42).

In this study, we sought to ensure that examples from uniquely functioning HLA groups were represented in our list of alleles to analyze, even if they were not present in majority populations. To achieve this, we needed to characterize functional similarity relationships between all HLAs in a way that would estimate the magnitude of difference between distinct HLAs and enable clustering at resolutions that vary from the curated nomenclature. Prior efforts have constructed HLA-I clusters and hierarchies based on distances defined by binding peptide set overlap or motif similarity identified from peptide binding data (2, 43). However, by the nature of their methods, such studies were limited to HLAs for which abundant data was available at the time and thus did not cover the full set we wished to consider.

To leverage all data used to train our binding prediction systems, we utilized a fixed length embedding of the full HLA amino acid sequence generated at an intermediate step in our trained neural networks as a basis for calculating distances between HLAs. For robustness, the embeddings from each model in the ensemble were stacked into one vector, and all clustering operations were performed on these concatenated ensemble embeddings. This approach to HLA grouping enabled consideration of all HLAs whose full sequence was known to our system (all data from the IPD-IMGT/HLA Database up to July 2019 (44, 45), including those for which limited empirical binding data is currently available.

To capture both local and global structure in high dimensional data, PHATE (46) was used to approximate relationships between our HLA embeddings in two dimensions (*knn* = 20 was used to focus more on global relations). For additional robustness to noise in cluster assignment, agglomerative clustering was used in the simplified PHATE coordinate space to group HLAs. Using a sweeping threshold across agglomerative clustering runs, a stable point was identified at 38 HLA-I clusters that maximized the silhouette score (**Fig. 13A**).

**Figure 13.**
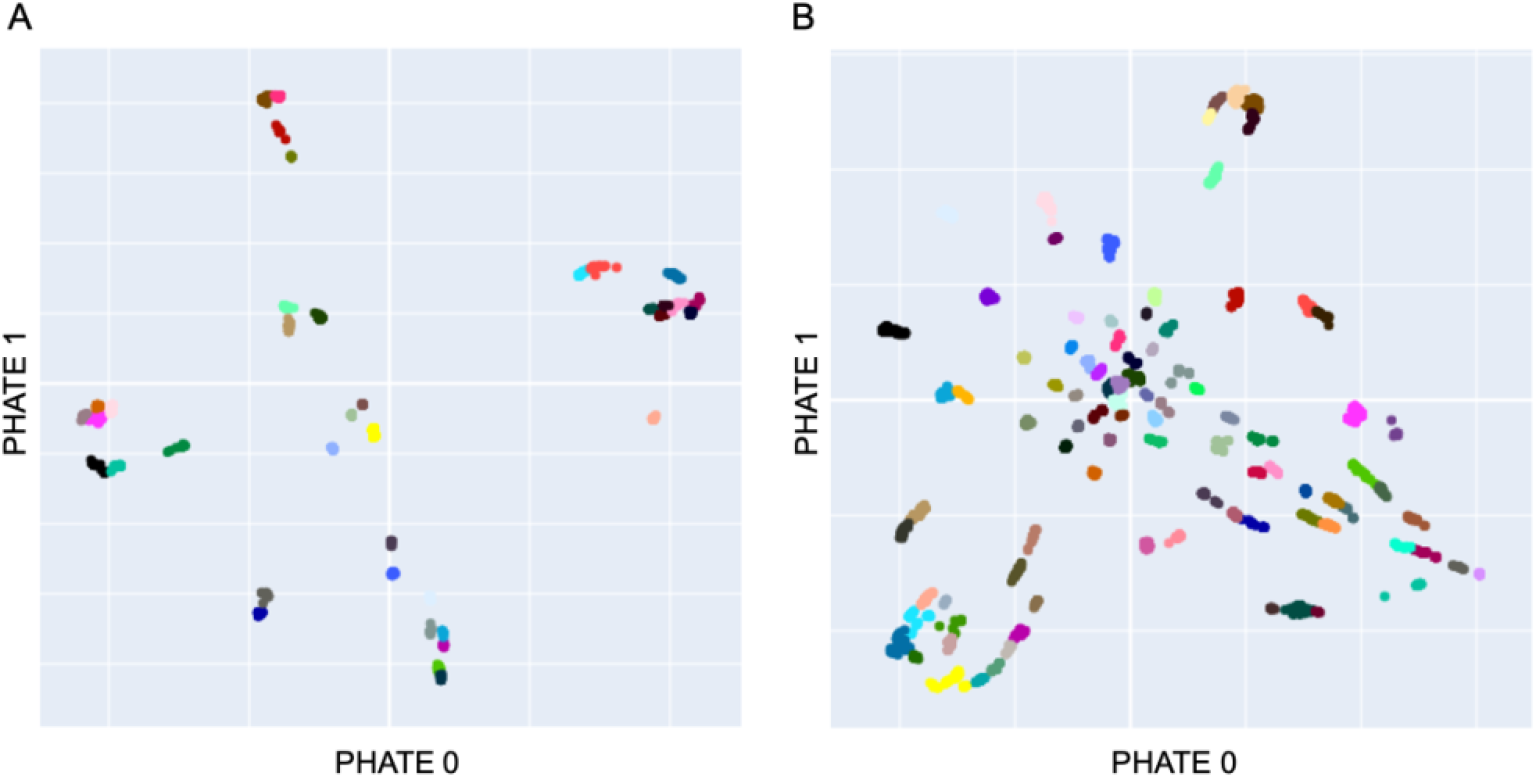
PHATE (46) visualization of HLA-I and HLA-II molecules. (A) Visualization of clustered RNN embeddings of all HLA-I A, B, and C molecules for which full amino acid sequences were available. Agglomerative clustering was used to group embeddings into 38 clusters, illustrated in the plot as discrete colors. (B) PHATE visualization and clustering of all permutations of compatible HLA-II alpha and beta chain embeddings, with a stable and silhouette score optimal point at 93 clusters.

Cluster assignments were then paired with data from the Allele Frequency Net Database (AFND) (47, 48) to select the set of HLA-I molecules for consideration in analysis. To first ensure all functional clusters were represented, the set was initially populated by the top 2 highest frequency HLAs from each of the 38 clusters. Subsequently, all HLAs with *f_HLA_* ≥ *min*(*μ,M*) – 0.6 * *σ* were added to the set (where *f_HLA_* is the frequency of appearance of an HLA, and *μ, M, σ* are the mean, median and standard deviation of the initial selection of 76 HLA-Is). The 0.6 multiplier was used so that the threshold ensured all top 100 most frequent HLA-I molecules were included. The final selection of 114 HLA-I molecules thus spanned all functional groups at the resolution of our clusters as well as the highest frequency alleles.

### HLA-II grouping and selection

The same studies that had previously approached HLA-I functional grouping (2, 43) did not attempt clustering of HLA-II. Using our trained neural nets for predicting HLA-II binding, we followed a similar pipeline as with HLA-I for creating functional clusters and selecting a comprehensive set for analysis.

In the case of HLA-II, our neural network architectures first process the full amino acid sequence of the alpha and beta chains independently, before allowing the information to be combined for binding prediction. These independent intermediate representations constitute the embeddings used to represent HLA-II alpha and beta chains for clustering. As with HLA-I, each representation was a stack of embeddings from all 5 models in an ensemble.

HLA-II clusters were assigned at 3 resolutions. For alpha chains alone, we found 18 clusters via agglomerative clustering and threshold sweep; and 16 clusters for all beta chains considered. Note that only alpha and beta chains with full amino acid sequences available (IPD-IMGT/HLA Database up to July 2019 (44, 45) were used. Finally, we considered the space of all compatible permutations of alpha + beta pairs. Each compatible pair (for example: DQA1*05:01-DQB1*06:02) was represented by the concatenation of the independent alpha and beta embedding, and all such pairs were represented in two dimensions with PHATE (at *knn* = 20). A total of 93 clusters were assigned to explain all alpha + beta permutations with our agglomerative approach (**Fig. 13B**). The full lists of HLA-I and HLA-II cluster membership assignments are provided as Supplementary CSV files and interactive plots.

Selecting a comprehensive HLA-II set for analysis utilized all 3 clustering assignments to ensure a fair sampling. The top 2 highest frequency alpha chains were selected from each of the 18 alpha clusters (4 singular clusters only contributed 1 alpha chain), and all other alpha chains that satisfied *f_α_* ≥ *min*(*μ,M*) – 0.5 * *σ* were added to the set (where is the frequency of appearance of an HLA-II alpha chain, and *μ*, *M*, *σ* are the mean, median and standard deviation of the initial set of 32 alpha chains).

Independently, the top 3 highest frequency beta chains from each of the 16 beta clusters were selected, and supplemented with all other beta chains that met *f_β_* ≥ *min*(*μ,M*) – 0.5 * *σ* (where *f_β_* is the frequency of appearance of an HLA-II beta chain, and *μ*, *M*, *σ* are the mean, median and standard deviation of the initial set of 47 beta chains).

The above sets were then combined to create 645 compatible permutations of alpha + beta chain pairs. The top 2 HLA-II alpha + beta pairs from any of the 93 clusters that were not covered by the selected set were added; bringing the augmented total to 661 HLA-II alpha + beta pairs. Since haplotype information in the AFND (47, 48) was less comprehensive than allele frequencies for independent alpha and beta chains, pairs within clusters were sorted by *max*(*f_α_*, *f_β_*); first at 4 digit allele nomenclature resolution and secondarily at 2 digit allele nomenclature resolution.

Finally, a pruning stage removed some HLA-II alpha + beta pairs from each of the 93 clusters in the analysis set only if: they were not in the top 2 samples representing a unique combination of alpha cluster (18 labels) and beta cluster (16 labels), and *f_β_* ≤ 0.05 at the 4-digit HLA-II nomenclature resolution. This ensured that we kept all functionally unique HLA-II alpha + beta combinations and spanned the entire set of 93 clusters in our analysis set, but did not heavily oversample functionally similar pairs unless they appeared with very high frequency in the world population.

The final HLA-II set for analysis included 373 alpha + beta pairs, with 37 unique alpha chains and 66 unique beta chains represented. See **Supplementary Files** for details.

### HLA-I potential epitope hotspot localization

Protein regions that featured an increased frequency of peptides with high predicted HLA binding confidence relative to elsewhere on the same protein were identified as *potential epitope hotspots*. These hotspots helped narrow our search space, added robustness to noise for global comparisons, and enhanced the interpretability of global analysis by enabling localization of regions at most risk for potential epitope loss.

To detect hotspots, first an aggregate signal was independently obtained for each HLA-I in our analysis set. All peptides of lengths from 8 to 12 were considered in a sliding window fashion, and the value at each protein position was the average binding prediction of all overlapping peptides. **Figure 14A** illustrates these aggregate signals for the reference S protein of SARS-CoV-2 across all 114 HLA-I (rows).

**Figure 14.**
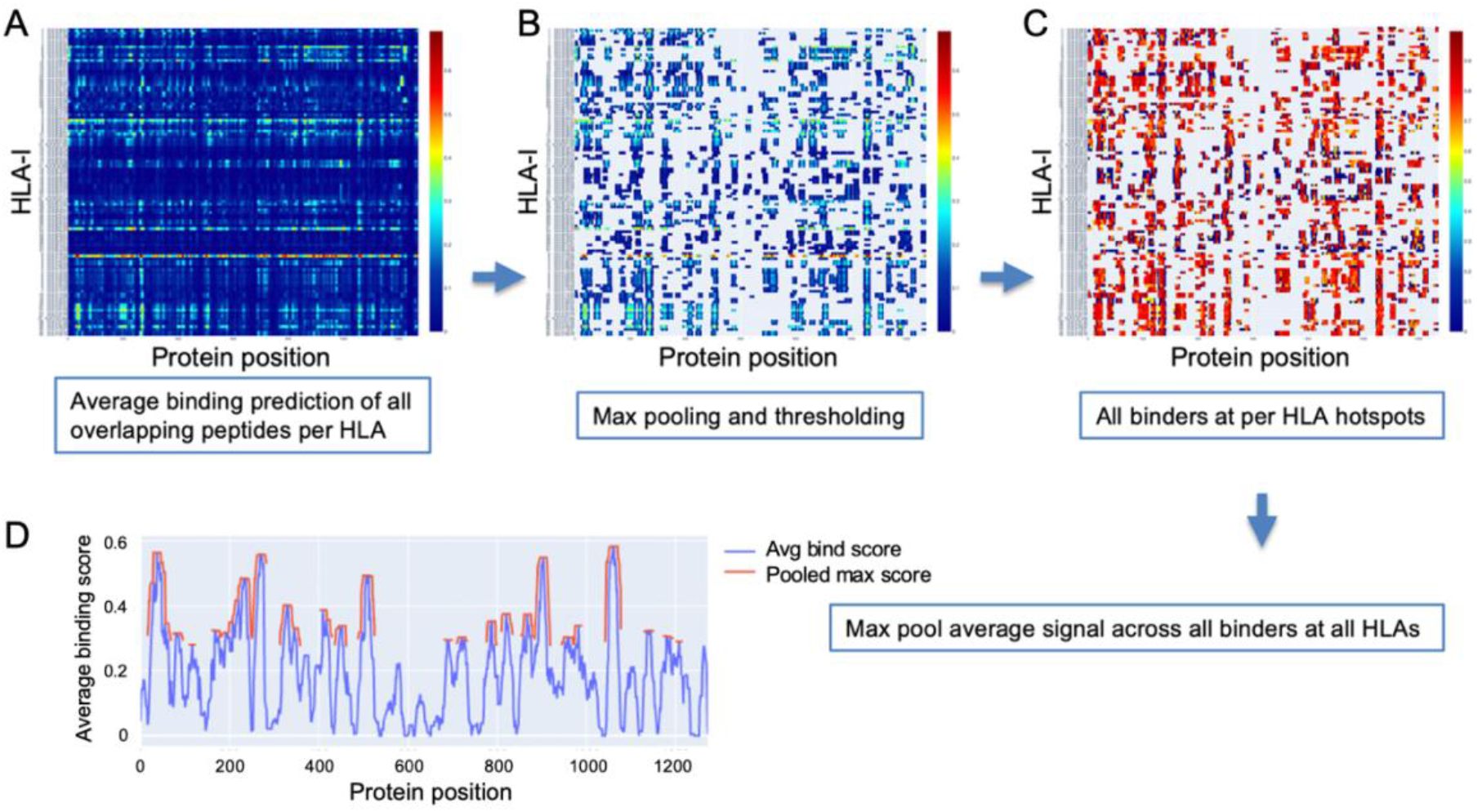
Method for detection of binder hotspots. (A) Binding prediction scores of all overlapping peptides at each position along the S protein (x-axis), averaged independently for each HLA-I molecule (y-axis). (B) Top binding score locations based on thresholding max pooled averaged prediction scores independently for each HLA (per-HLA hotspots). Unselected regions are masked out and appear in gray. Values before pooling are shown to keep relation to the prior plot clear. (C) Averaged scores of all peptides classified as binders that overlap per-HLA hotspots. (D) The final HLA-I hotspot locations (red) selected by max pooling and thresholding the masked binder signal averaged across HLAs (blue).

Max pooling with a window size based on the dominant binding peptide length (9 amino acids) was then applied to the aggregate signal for each HLA, and all values in the top 10% of maxima were selected as hotspots (**Fig. 14B**). The max thresholding operation was applied independently per HLA to help discount remaining uncorrected biases potentially causing systematic shift in prediction scores for some HLAs (ensuring peak binder locations of less well characterized HLAs were not discarded in later analysis).

Once potential epitope hotspots were selected independently for each HLA-I, they were used to narrow the scope of the aggregate signal to average only peptides classified as binders that overlap their respective HLA-I hotspots (**Fig. 14C**). These masked binder signals were averaged across HLAs to create a global HLA-I average binder score, which was then max pooled (size = 9), and the top 25% of maxima were selected as general HLA-I binding hotspots for the protein of interest (**Fig. 14D**).

### HLA-II potential epitope hotspot localization

The same procedure was followed for identification of HLA-II potential epitope hotspots as for HLA-I, except all peptide sizes considered ranged from 11 to 21, and all max pooling steps utilized the most frequent HLA-II ligand length of 15 (instead of 9 used in HLA-I). The set of selected HLA-II for analysis was also considerably larger: 373 alpha + beta pairs.

### Pan-HLA candidate epitope hotspot localization

To identify locations along a protein with the highest concentration of peptides predicted to bind both HLA-I and HLA-II with high confidence, we averaged the global HLA-I and HLA-II signals before their respective final max pooling steps were applied. This pan-HLA average binder signal was then max pooled (with size = 12) and hotspot locations were selected based on the top 25% of the max pooled signal. **Figure 1** illustrates HLA-I, HLA-II, and pan-HLA hotspot locations for 6 key proteins from the SARS-CoV-2 reference genome.

### SARS-CoV-2 genome data

SARS-CoV-2 genome and protein data were downloaded from the NIH NCBI portals (https://www.ncbi.nlm.nih.gov/datasets/coronavirus/genomes/, https://www.ncbi.nlm.nih.gov/datasets/coronavirus/proteins/) respectively on the dates of March 19, May 11, July 27, September 25, and November 26, 2021. Upon download all genomes were classified with the latest version of the Pangolin tool available at each time point (49), to enable analysis with respect to the PANGO lineage classifications (50).

To base our viral surveillance on the most confident and critical data, our analyses of the S and N proteins considered only samples for which the S or N protein sequences were observed at least 3 times (**Fig. 4**).

### Potential epitope hotspot alignment across variants

Potential epitope hotspots for all considered proteins were localized based on the SARS-CoV-2 reference genome (NCBI Reference Sequence: NC_045512 (23). To map hotspot locations onto each new mutated version of a protein as it was processed, dynamic time warping (DTW) (51) was used. This enabled direct comparison of relative predicted HLA binder counts at hotspot locations between viral variants.

### Ranking protein variants by sum fraction of binders lost

To summarize how mutations in a protein variant, *p*, impact its predicted peptide binding, we defined the *binder count fraction* for each HLA molecule, *m*, as:

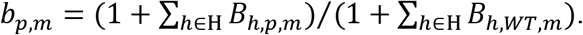

*B_h,p,m_* is the number of peptides at the hotspot, *h*, in protein variant, *p*, that were classified as binders for a specific HLA, *m; B_h,WT,m_* is the corresponding binder count in the wildtype (WT) version of the protein being considered; and *H* is the set of all corresponding hotspot regions in *WT* and *p*. All wildtype proteins sequences were obtained from the SARS-CoV-2 reference genome (NCBI Reference Sequence: NC_045512).

The overall potential for epitope loss of a protein variant when compared to wild type was captured by the *sum fraction of binders lost*, defined as:

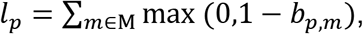

where M is our set of all HLA molecules selected for analysis. Cases where epitope count for a specific HLA increased compared to the reference were intentionally omitted from the summation to focus only on the risk of losing epitopes that may be critical to immune response.

To compare overall binder loss magnitudes across time and across class I and class II, *sum fraction of binders lost* was divided by the number of HLAs considered to obtain *average bind loss per HLA*.

Independently for HLA-I and HLA-II, all versions of the S and N protein at each data time point were ranked according to *sum fraction of binders lost*, such that protein versions with the most significant overall loss in HLA binders appeared at the top. Prior ranking and all later stages of analysis, any protein version that featured unspecified amino acids with a quantity greater than 1% of protein length were removed from consideration. A fractional value (*bind loss rank fraction*) was assigned to represent ranking position; where values near 0 correspond to the top of the ranking order (most loss of HLA binders), and values near 1 indicate a rank at the end (least loss of HLA binders). *Bind loss rank fraction* was used in the analysis illustrated in **Figure 6** to select lineages whose representative S ranked within a percentage of unique proteins exhibiting most binder loss, as captured by the plot’s x-axis.

### Capturing the emergence of B. 1.1.529 with supplementary data from GISAID

To supplement NIH NCBI data from November 26, 2021 to cover the emergence and rapid spread of the B.1.1.529 (Omicron) lineage, we obtained all Omicron genomes from GISAID (29–31) up to December 6, 2021. Genomes were required to be complete, and those with low coverage were excluded from the query. Each downloaded genome was aligned to reference using MAFFT (52). All unique versions of S and N proteins that included less than 1% unspecified amino acids and were classified by GISAID as Omicron were combined with the November 26 data. This augmented December 6 dataset was then used to capture a more timely picture of relative potential for epitope loss across VOC lineages. To verify that our early Omicron data remained representative of the rapidly spreading lineage, we used two additional time points (December 20, 2021 and December 29, 2021) to confirm that the top most frequent versions of S and N proteins representing the lineage had in fact not changed. Thus, our analysis remained representative throughout the remainder of 2021.

### Characterizing binding loss variation at pan-HLA hotspots across lineages

The B.1.1.529 augmented December 6, 2021 data was used to analyze the full range of epitope loss potential in VOC and VOI lineages at all pan-HLA hotspot regions (**Fig. 9**). To summarize changes, binder count fraction relative to reference (WT) was first computed independently per hotspot, *h*, then averaged across the set of all HLAs, *M*, for each protein, *P*: *b_h,p_* = ∑*m*∈M((1 + *B_h,p,m_*)/(1 + *B_h,WT,m_*))/|M|.

For each VOC and VOI lineage, box plots in **Figure 9** were created by collecting *b_h,p_* values of all unique protein sequence instances that included at least one corresponding source genome classified as the lineage being considered. Finally, *b_h,p_* values (averaged across HLA-I) for all most frequent instances of S and N proteins representing all VOC lineages are illustrated in **Figure 13**.

## References

1. J. Prévost, A. Finzi, The great escape? SARS-CoV-2 variants evading neutralizing responses. Cell Host Microbe 29, 322–324 (2021).

2. J. Sidney, B. Peters, N. Frahm, C. Brander, A. Sette, HLA class I supertypes: a revised and updated classification. BMC Immunol 9, 1 (2008).

3. C. I. Stoddard et al., Epitope profiling reveals binding signatures of SARS-CoV-2 immune response in natural infection and cross-reactivity with endemic human CoVs. Cell Rep 35, 109164 (2021).

4. C. R. Moderbacher et al., Antigen-specific adaptive immunity to SARS-CoV-2 in acute COVID-19 and associations with age and disease severity. Cell 183, 996–1012 (2020).

5. T. Sekine et al., Robust T cell immunity in convalescent individuals with asymptomatic or mild COVID-19. Cell 183, 158 (2020).

6. A. T. Tan et al., Early induction of functional SARS-CoV-2-specific T cells associates with rapid viral clearance and mild disease in COVID-19 patients. Cell Rep 34, 108728 (2021).

7. A. Sette, S. Crotty, Adaptive immunity to SARS-CoV-2 and COVID-19. Cell 184, 861–880 (2021).

8. A. Bertoletti, N. Le Bert, M. Qui, A. T. Tan, SARS-CoV-2-specific T cells in infection and vaccination. Cell Mol Immunol 18, 2307–2312 (2021).

9. A. Grifoni et al., SARS-CoV-2 human T cell epitopes: Adaptive immune response against COVID-19. Cell Host Microbe 29, 1076–1092 (2021).

10. J. Heide et al., Broadly directed SARS-CoV-2-specific CD4+ T cell response includes frequently detected peptide specificities within the membrane and nucleoprotein in patients with acute and resolved COVID-19. PLoS Pathog 17, e1009842 (2021).

11. O. W. Ng et al., Memory T cell responses targeting the SARS coronavirus persist up to 11 years post-infection. Vaccine 34, 2008–2014 (2016).

12. N. Le Bert et al., SARS-CoV-2-specific T cell immunity in cases of COVID-19 and SARS, and uninfected controls. Nature 584, 457–462 (2020).

13. A. Grifoni et al., Targets of T Cell Responses to SARS-CoV-2 Coronavirus in Humans with COVID-19 Disease and Unexposed Individuals. Cell 181, 1489 (2020).

14. A. Tarke et al., Comprehensive analysis of T cell immunodominance and immunoprevalence of SARS-CoV-2 epitopes in COVID-19 cases. Cell Rep Med 2, 100204 (2021).

15. S. K. Saini et al., SARS-CoV-2 genome-wide T cell epitope mapping reveals immunodominance and substantial CD8(+) T cell activation in COVID-19 patients. Sci Immunol 6 (2021).

16. A. Tarke et al., Impact of SARS-CoV-2 variants on the total CD4+ and CD8+T-cell reactivity in infected or vaccinated individuals. Cell Reports Medicine 2 (2021).

17. C. Lucas et al., Impact of circulating SARS-CoV-2 variants on mRNA vaccine-induced immunity. Nature 10.1038/s41586-021-04085-y (2021).

18. B. Agerer et al., SARS-CoV-2 mutations in MHC-I-restricted epitopes evade CD8(+) T cell responses. Sci Immunol 6 (2021).

19. R. Khandia et al., Emergence of SARS-CoV-2 Omicron (B.1.1.529) variant, salient features, high global health concerns and strategies to counter it amid ongoing COVID-19 pandemic. Environ Res 209, 112816 (2022).

20. E. Gabitzsch et al., Dual-Antigen COVID-19 Vaccine Subcutaneous Prime Delivery With Oral Boosts Protects NHP Against SARS-CoV-2 Challenge. Front Immunol 12, 10.3389/fimmu.2021.729837 (2021).

21. F. Krammer, SARS-CoV-2 vaccines in development. Nature 586, 516–527 (2020).

22. P. Sieling et al., Prime hAd5 Spike plus Nucleocapsid Vaccination Induces Ten-Fold Increases in Mean T-Cell Responses in Phase 1 Subjects that are Sustained Against Spike Variants. medRxiv 2021.04.05.21254940, doi: 10.1101/2021.1104.1105.21254940 (2021).

23. GenBank, NCBI Reference Sequence NC_045512.2. GeBank https://www.ncbi.nlm.nih.gov/nuccore/NC_045512 (2020).

24. J. Lan et al., Structure of the SARS-CoV-2 spike receptor-binding domain bound to the ACE2 receptor. Nature 581, 215–220 (2020).

25. CDC, SARS-CoV-2 Variant Classifications and Definitions. Centers for Disease Control and Prevention (CDC) Website, https://www.cdc.gov/coronavirus/2019-ncov/variants/variant-info.html#unweighted-proportions-substitutions-of-therepeutic-concern (2021).

26. S. K. Dhanda et al., Development of a novel clustering tool for linear peptide sequences. Immunology 155, 331–345 (2018).

27. S. K. Dhanda et al., ImmunomeBrowser: a tool to aggregate and visualize complex and heterogeneous epitopes in reference proteins. Bioinformatics 34, 3931–3933 (2018).

28. R. Vita et al., The Immune Epitope Database (IEDB): 2018 update. Nucleic Acids Res 47, D339–D343 (2019).

29. S. Khare et al., GISAID’s Role in Pandemic Response. China CDC Wkly 3, 1049–1051 (2021).

30. S. Elbe, G. Buckland-Merrett, Data, disease and diplomacy: GISAID’s innovative contribution to global health. Global challenges (Hoboken, NJ) 1, 33–46 (2017).

31. Y. Shu, J. McCauley, GISAID: Global initiative on sharing all influenza data - from vision to reality. Euro Surveill 22, 30494 (2017).

32. A. Nguyen et al., Human Leukocyte Antigen Susceptibility Map for Severe Acute Respiratory Syndrome Coronavirus 2. Journal of virology 94 (2020).

33. T. J. O’Donnell et al., MHCflurry: Open-Source Class I MHC Binding Affinity Prediction. Cell Syst 7, 129–132.e124 (2018).

34. B. Reynisson, B. Alvarez, S. Paul, B. Peters, M. Nielsen, NetMHCpan-4.1 and NetMHCIIpan-4.0: improved predictions of MHC antigen presentation by concurrent motif deconvolution and integration of MS MHC eluted ligand data. Nucleic Acids Res 48, W449–w454 (2020).

35. V. Jurtz et al., NetMHCpan-4.0: Improved Peptide-MHC Class I Interaction Predictions Integrating Eluted Ligand and Peptide Binding Affinity Data. Journal of immunology (Baltimore, Md.: 1950) 199, 3360–3368 (2017).

36. H. Pearson et al., MHC class I-associated peptides derive from selective regions of the human genome. J Clin Invest 126, 4690–4701 (2016).

37. Y. Cui, M. Jia, T.-Y. Lin, Y. Song, S. Belongie, Class-Balanced Loss Based on Effective Number of Samples. CVPR 2019 arXiv: 1901.05555 [cs.CV] (2019).

38. D. Gfeller et al., The Length Distribution and Multiple Specificity of Naturally Presented HLA-I Ligands. Journal of immunology (Baltimore, Md.: 1950) 201, 3705–3716 (2018).

39. M. Bassani-Sternberg et al., Deciphering HLA-I motifs across HLA peptidomes improves neo-antigen predictions and identifies allostery regulating HLA specificity. PLoS Comput Biol 13, e1005725 (2017).

40. B. Lakshminarayanan, A. Pritzel, B. C., Simple and Scalable Predictive Uncertainty Estimation using Deep Ensembles. 31st Conference on Neural Information Processing Systems (NIPS) https://papers.nips.cc/paper/2017/file/9ef2ed4b7fd2c810847ffa5fa85bce38-Paper.pdf (2017).

41. WHO, Nomenclature for factors of the HL-a system. Bull World Health Organ 39, 483–486 (1968).

42. S. G. E. Marsh et al., Nomenclature for factors of the HLA system, 2010. Tissue Antigens 75, 291–455 (2010).

43. D. Gfeller, M. Bassani-Sternberg, Predicting Antigen Presentation-What Could We Learn From a Million Peptides? Front Immunol 9, 1716 (2018).

44. IPD-IMGT/HLA, IPD-IMGT/HLA Database. www.ebi.ac.uk/pd/imgt/hla/ (2021).

45. J. Robinson et al., IPD-IMGT/HLA Database. Nucleic Acids Res 48, D948–d955 (2020).

46. K. R. Moon et al., Visualizing structure and transitions in high-dimensional biological data. Nat Biotechnol 37, 1482–1492 (2019).

47. F. F. Gonzalez-Galarza et al., Allele frequency net database (AFND) 2020 update: gold-standard data classification, open access genotype data and new query tools. Nucleic Acids Res 48, D783–d788 (2020).

48. T. A. F. N. Database, Allele Frequency Net Database (AFND). www.allelefrequencies.net (2021).

49. Á. O’Toole et al., Assignment of epidemiological lineages in an emerging pandemic using the pangolin tool. Virus Evolution 10.1093/ve/veab064 (2021).

50. A. Rambaut et al., A dynamic nomenclature proposal for SARS-CoV-2 lineages to assist genomic epidemiology. Nature Microbiology 5, 1403–1407 (2020).

51. T. Giorgino, Computing and Visualing Dynamic Time Warping Alignments in R: The dtw Package. Journal of Statistical Software 31, i07 (2009).

52. K. Katoh, D. M. Standley, MAFFT multiple sequence alignment software version 7: improvements in performance and usability. Mol Biol Evol 30, 772–780 (2013).

